# Evaluating Biodiversity Credit Metrics Using Metacommunity Modelling

**DOI:** 10.1101/2024.06.03.597228

**Authors:** Dominik M Maczik, Vincent A.A. Jansen, Axel G. Rossberg

## Abstract

Global biodiversity enhancement is central to the UN Sustainable Development Goals and climate change mitigation. Achieving the Kunming-Montreal Global Biodiversity Framework’s ‘30 by 30’ target requires an estimated additional US$700 billion annually. Biodiversity credit markets seek to address this funding gap by assigning financial value to biodiversity and ecosystem services. However, limited understanding of the metrics underpinning these credits pose significant barriers to their scalability and effectiveness.

This pioneering study compares six credit metrics with six established biodiversity metrics to assess whether methodology choice influences metric responses to different ecosystem perturbations, identifying metrics best suited for specific interventions, and exploring comparability across metrics. A spatially explicit, multi-layered metacommunity simulation model, capable of reproducing a variety of empirically established macro-ecological patterns, was adapted to track ecosystem responses to six perturbation experiments and to record changes in the twelve tracked biodiversity metrics.

Results reveal substantial divergence in how credit metrics assign value to nature, particularly between those estimating ecosystem services and those assessing species extinction risk. These findings underscore the need for careful alignment between metric selection and the ecological objectives of biodiversity projects and suggest that the development of a universal biodiversity credit is unlikely. Furthermore, in addition to metrics estimating ecosystem services, our results suggest that projects should incorporate metrics that are sensitive to declines in species-level abundances, thereby reflecting extinction risk.

## 1. Introduction

Biodiversity is in a global, anthropogenic decline, primarily driven by economic activity, widely considered to be the sixth mass extinction (Ceballos et al., 2015; Díaz et al., 2019; Jaureguiberry et al., 2022). Concurrently, studies suggest that over half of global GDP, approximately $40 trillion, exhibits moderate to high dependency on nature and ecosystem services (Dasgupta, 2021; IPBES, 2019). Resource constraints have consistently been identified as a significant barrier to achieving biodiversity targets in the literature (Deutz et al., 2020; Seidl et al., 2024; Xu et al., 2021). Current public investment in biodiversity conservation is only an approximate $121 billion, representing merely 0.19-0.25% of global GDP, while econometric modelling suggests an annual global finance gap of $598-824 billion (Deutz et al., 2020; Seidl et al., 2020). Over the past decades, private conservation finance mechanisms have not been of sufficient scale to address the biodiversity funding gap (Dempsey & Suarez, 2016; Deutz et al., 2020). Consequently, beyond merely addressing the funding gap, transformative change is necessary to tackle the underlying drivers of biodiversity loss (Dempsey et al., 2022; McElwee et al., 2020; Otero et al., 2020).

The Convention on Biological Diversity (CBD) defines biodiversity as complexity arising at the genetic, species and ecosystem levels of community organization. In addition, Target 2 of the Kunming-Montreal Global Biodiversity Framework calls for the restoration of 30% of global degraded ecosystems, “in order to enhance biodiversity and ecosystem functions and services, ecological integrity” (CBD, 2022). The integration of these diverse elements into unified metrics presents significant challenges, resulting in the proposal of various metrics and certificates. Indeed, adequate quantification and measurement of biodiversity change remain fundamental obstacles in the assessment of the efficacy of conservation and restoration projects (Damiens et al., 2021; Karolyi & Tobin-de la Puente, 2023; Needham et al., 2019). We have insufficient understanding not only of the characteristics of the metrics proposed for the quantification of biodiversity, but also of what component of biodiversity they measure.

‘Biodiversity credit’ has emerged as a catch-all term representing a wide variety of certificates for evidence-based units of positive biodiversity outcomes (Biodiversity Credit Alliance, 2024). For clarity, this paper refers to these financial instruments as “credits”, the metrics that credit schemes utilise to measure biodiversity as “credit metrics”, while the term “metric” may refer to any quantitative measure. The existing biodiversity credit metric landscape is marked by significant fragmentation (Kim et al., 2025). Many credit metrics aim for a broad assessment of biodiversity, while others focus on specific aspects of conservation and restoration, for instance extinction risk (de Azevedo-Gonçalves & Gonçalves, 2024). They may be calculated using a combination of species abundance data and habitat and ecosystem characteristics, with one analysis suggesting that over half of credit metrics studied rely on a combination of species and ecosystem data (Wunder et al., 2025). To date it is not clear what impact differences in credit methodologies have in practice, and therefore which methodologies should be favoured.

One might propose that the eventual market success of a handful of biodiversity credit schemes will reflect the result of natural societal consensus building on the correct valuation of nature. However, that would allow humanity’s relationship with nature to be solely defined through market dynamics, which can be morally problematic, subject to market failures, and may lead to inefficient use of conservation potential (Cinner et al., 2021; Lockie, 2019; Miles et al., 2025; Neuteleers, 2022). Indeed, one might argue that the appropriate valuation of nature and ecosystems presents a complex challenge that defies universal standardisation, as these assessments are inherently contingent upon local scientific, socioeconomic, and cultural considerations (Jones et al., 2019). On the other hand, due to the globally interconnected nature of ecosystems as well as corporate value chains, a unifying global perspective is also required. The existence of these two competing lines of reasoning is particularly evident in biodiversity credit schemes, where the absence of standardised metrics reflects the fundamental challenge of translating valuations of nature into quantifiable units. Any misalignment between credit metrics and progress towards such societally agreed global positive end states or local needs can lead to undervaluing the achieved results, or underdelivering on the promised environmental objectives. An important step in preventing such misalignment is to understand how different metrics respond to different forms of biodiversity change.

To investigate this, this paper addresses the following key questions that users of biodiversity credits want to know: (1) do all credits respond in a similar fashion to diverse forms nature perturbations; (2) what kind of projects are suitable to generate credits measured in a particular way or, if the project is decided based on other considerations, which metrics are suitable to quantify attainment of the projects’ biodiversity objectives; (3) if and how credits based on different metrics can be compared; and (4) whether generation of credits actually contributes to globally agreed objectives. To address these questions, we investigated the responses of twelve metrics to six simulated nature perturbation experiments representing societal interventions in nature. Six of these metrics formed the basis of biodiversity crediting schemes, while the other six are ecological metrics used to quantify progress in achieving globally agreed objectives.

## 2. Methods

### 2.1. The Lotka-Volterra Metacommunity Model

By describing a group of local ecological communities interlinked by dispersal, metacommunity models permit simulation of landscape-level ecological dynamics. To understand the responses of credit metrics to interventions, we performed computer simulations of complex metacommunities using the Lotka-Volterra Metacommunity Model (LVMCM) (O’Sullivan et al., 2019). This established, spatially explicit metacommunity model reproduces a large variety of empirically well-documented macro-ecological patterns, including those seen in occupancy frequency distributions, range size distribution and other aspects of the distribution of species across space and time (O’Sullivan et al., 2021, 2023). The model combines multiple elements considered essential for metacommunity dynamics. The base layer describes the spatial variation of environmental variables, thus representing nature’s diverse habitats. For the purposes of this study only two environmental variables were used which were sampled from a spatially correlated 2D random field with the mean fixed at 0. For each environmental variable, each simulated species is assigned an optimal value that defines the species’ preferred abiotic conditions. The species’ intrinsic growth rates are based on this optimal value and the niche width, which is kept uniform across all species as a simplifying assumption. N= 10000 simulations had mean growth rate values at 1 per unit time, while standard deviation at 0.70363 = 0.4951^1/2^ per unit time. The simulation area is subdivided into a 10 by 10 lattice of sites, and adjacent sites are connected through dispersal. A network of adverse ecological interactions between the species, identical across sites, is constructed by random sampling. These interactions introduce an element of competition and thus limit the number of species that can co-exist at a given site. Further information on the Lotka-Volterra competition models governing species behaviour is available in Supporting Information.

The simulation protocol begins by sampling the spatially autocorrelated environmental variables. This is followed by the assembly of an ecological community by iterated invasion of species with randomly generated traits (environmental preferences and species interactions). This process simulates the natural assembly of a real-world ecosystem until species richness fluctuates around a steady state value. O’Sullivan et al. (2019) have shown that such assembly is required to reproduce realistic macroecological patterns in the LVMCM. We therefore use it here to model pristine and undisturbed ecosystems. We assembled n=10000 of such pristine communities which contained between 23 and 99 species (53.28 on average), with an average range of 35 out of 100 sites. After the simulated experimental intervention, as described below, population dynamics are simulated over a period of ten unit times, where one unit time corresponds to the typical time required for total population biomass to reach a new equilibrium (Figure 8). This choice represents a balance between the potentially long response times of complex metacommunities (O’Sullivan et al., 2023), and the much shorter times to credit issuance required for efficient markets. The resulting community dynamics are modelled while the size and geographical distribution of species populations are measured to calculate biodiversity metrics, including credit metrics, and their change over time.

Within the LVMCM, a species is considered locally extinct at a given site if its biomass falls below a predetermined extinction threshold and is considered globally extinct if it is extinct in all sites. In this study, because our simulated metacommunities consisting of only 100 linked sites are small compared to typical continental communities, we anticipate our simulations to overestimate the risks of primary and secondary extinctions resulting from interventions into the ecosystem.

### 2.2. Metric and credit selection

Using the LVMCM, we simulated six different nature restoration scenarios and quantified the resulting change using twelve different metrics, six of which form the basis of biodiversity crediting schemes. In addition, to support contextualisation of the credit metrics modelled, several established ecological metrics are evaluated by adapting published methodologies to the kind of data generated by the LVMCM (abundances and spatial distributions of a group of competing species). These are the overall population biomass, species richness, Range Size Rarity for each site, the Living Planet Index (Collen et al., 2009), Mean Species Abundance (Alkemade et al., 2009), and the Species Threat Abatement and Restoration

Metric (STARt) (Mair et al., 2021). Certain metrics rely on additional locally or globally determined attributes, such as indicator species, the IUCN Red List, and the IUCN Red List of Ecosystems (Keith et al., 2013; Myers et al., 2000; Rodrigues et al., 2006). The IUCN Red List categories were modelled based on species abundances only, with abundance thresholds fixed after assembly such as to match empirical proportions of species in Red List categories. STARt and RSR were calculated for the project area, while the biomass, extinction ratio, RLI, LPI and MSA are calculated globally, in line with their intended uses. A detailed description of how we modelled determination of these attributes is provided in Supporting Information.

Our selection of credit metrics followed a comprehensive review of documentations from 35 biodiversity credit issuers and nature-positive project developers, prioritising feasibility of implementation in our modelling framework. As biodiversity credits are a novel product in a nascent market, limited availability of clear and public documentation of methodology precluded the inclusion of some credit metrics in this thesis. The LVMCM tracks species in different environmental and spatial settings, which is why tracking metrics that use raw biological data is relatively straightforward. Other credit metrics rely on additional ecological data that is not straightforwardly modelled in the LVMCM. These include, e.g., genomic data, taxonomic relationships of species, eDNA, as well as expert judgements by field ecologists or other stakeholders. To evaluate such metrics other approaches will be required. Yet, the credit metrics we chose to model in this study cover a variety of fundamentally different methodologies.

The selected metrics are listed in Table 1. To maintain strict analytical neutrality and avoid any appearance of publicly endorsing or criticising specific credit schemes, we employ standardised identifiers for credit metrics throughout our analysis, as detailed in Table 1. All credit metrics were calculated for the project area, apart from *credit metric B*, which does not take area size into account. *Credit metrics B* and *D* rely on only species level data, *credit metrics A* and *F* require both species and habitat-based markers, and *credit metrics C* and *E* focus primarily on ecosystem-wide data. Where credit schemes require additional calculations, for instance to account for leakage or buffers, we decided not to include these calculations to maintain comparability of metrics. When credit schemes provide for flexibility in the implementation of metrics, the options with the least complexity were prioritised. This aligns with project developers’ incentives and therefore tests the credit metrics’ implementation in a realistic manner. Detailed information on the implementation of metrics is available in Supporting Information.

**Table 1.**
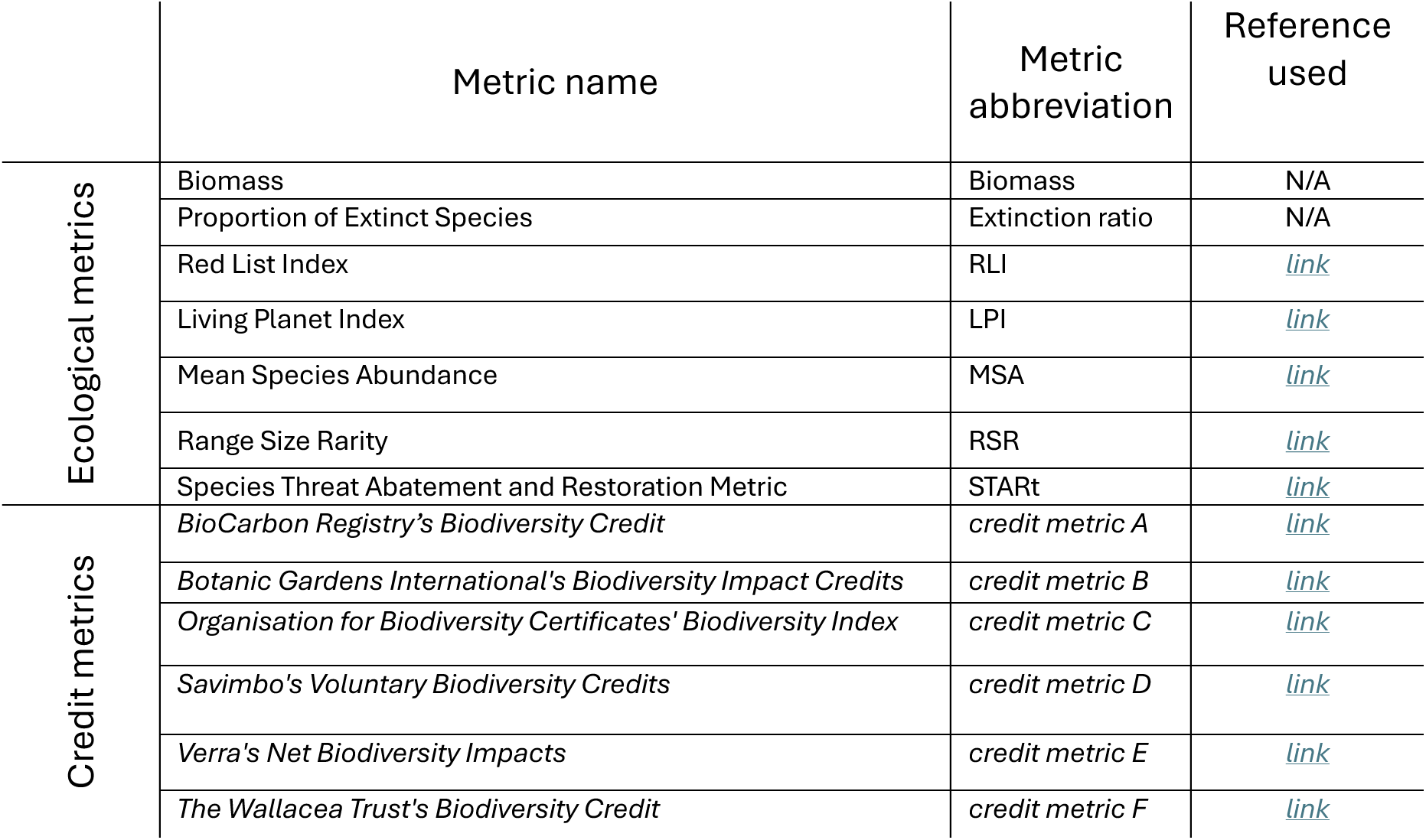
offers an overview of the twelve biodiversity metrics analysed. Six are established ecological metrics used to characterise community structure, and six are derived from biodiversity crediting schemes. Metrics are grouped by type along the y-axis, with name, abbreviation, and reference listed along the x-axis. Standardised identifiers are used for credit metrics to ensure analytical neutrality and avoid implicit endorsement or critique of specific schemes.

|  | Metric name | Metric abbreviation | Reference used |
| --- | --- | --- | --- |
| Ecological metrics | Biomass | Biomass | N/A |
|  | Proportion of Extinct Species | Extinction ratio | N/A |
|  | Red List Index | RLI | link |
|  | Living Planet Index | LPI | link |
|  | Mean Species Abundance | MSA | link |
|  | Range Size Rarity | RSR | link |
|  | Species Threat Abatement and Restoration Metric | STARt | link |
| Credit metrics | *BioCarbon Registry's Biodiversity Credit* | *credit metric A* | link |
|  | *Botanic Gardens International's Biodiversity Impact Credits* | *credit metric B* | link |
|  | *Organisation for Biodiversity Certificates' Biodiversity Index* | *credit metric C* | link |
|  | *Savimbo's Voluntary Biodiversity Credits* | *credit metric D* | link |
|  | *Verra's Net Biodiversity Impacts* | *credit metric E* | link |
|  | *The Wallacea Trust's Biodiversity Credit* | *credit metric F* | link |

### 2.3. Experimental protocol

The simulated experiments took place on the entire simulated field of 10 × 10 sites, except in the ‘*area increase*’ experiments explained below. The experiments were carried out using six kinds of modifications of the simulated environment, chosen to model a range common human interventions in nature. As detailed below, these are *environmental degradation, environmental restoration, (species-specific) extinctions, restoration area increase, species-specific restoration, and environmental heterogeneity restoration*.

#### 2.3.1. Environmental Degradation

To model general environmental degradation, the intrinsic growth rates of all species were incrementally reduced at every site. Initial, reductions in growth rate (adding -1 per unit time) lead to the gradual decline of most species, with the extinction of the most vulnerable. Larger decreases (adding -2) resulted in more significant extinction events, while extreme interventions (adding -3) caused the extinction of nearly all species and therefore a catastrophic collapse of the ecosystem. A control condition of simulating no environmental changes for ten unit times was conducted.

#### 2.3.2. Environmental Restoration

Environmental restoration was modelled by first modelling environmental degradation as above, removing species that went extinct, simulating ten unit times, and then restoring the original intrinsic growth rates of all remaining species. A control condition of simulating no environmental changes for ten unit times was conducted.

#### 2.3.3. Restoration Area Increase

Most biodiversity credit metrics, including 5 out of the 6 examined here, are calculated as the product of project area size and a factor computed from various biodiversity metrics. However, the emphasis on the project area varies between methodologies. To quantify the role of the project area compared to the surrounding buffer, we simulated the degradation – restoration sequence as above but modified intrinsic growth rates only in an *n* × *n* square (*n* = 1, 2, 3, 4) positioned as close to the centre of the 10 × 10 lattice as possible. To assess the methodologies’ impact, experimental results are compared against a control experiment where the same sized degraded environment is assessed over time but not restored. The time between degradation and restoration is again ten unit times. The metric calculations were restricted to the project area if the methodology provided for this. In fact, all metrics were calculated for the specified project area, except from *credit metric B*, biomass, the extinction ratio, LPI, MSA and RLI. A control condition of simulating no environmental changes at the given restoration area, for ten unit times was conducted.

#### 2.3.4. Extinction

Species-specific extinctions (e.g. due to disease of overharvesting) were modelled by removing 5, 10 or 15 randomly chosen species from the assembled community. Species’ intrinsic growth rates were left unchanged. Species removal frequently resulted in secondary and tertiary extinctions but also generated opportunities for the remaining species to occupy more patches. A control condition of simulating no changes for ten unit times was conducted.

#### 2.3.5. Species-specific Restoration

Field restoration efforts often focus on a single species that is threatened or of ecological or cultural significance (Lindenmayer et al., 2007; Runge et al., 2019; Simberloff, 1998). Here, we conduced numerical experiments where we investigated how metrics responded in the face of species-specific restoration efforts. In the LVMCM, after an initial saturation of the assembled community, the rarest species was targeted by progressively increasing its intrinsic growth rate by adding a constant between 0 and 0.2 in steps of 0.01. Small additions can benefit the entire community; however, with additions above a certain threshold the targeted species may adversely affect other species, resulting in an overall detrimental impact on the community. A control condition of simulating no changes for ten unit times was conducted.

#### 2.3.6. Environmental Heterogeneity Restoration

Ecologists and conservationist highlight the importance of environmental heterogeneity for the maintenance of biodiversity (Larkin et al., 2016; Rocchini et al., 2010; Stein et al., 2014). To model environmental heterogeneity restoration after homogenisation of the environment, we first set the random field representing all environmental variables to 0 for all sites and simulated relaxation of the metacommunity for ten unit times, removing extinct species. Restoration was then modelled by restoring the original values of the environmental variable and again simulating ten unit times. Experiments were conducted using one or two environmental variables, representing different levels of environmental complexity. A control condition of simulating no environmental changes for ten unit times was conducted.

### 2.4. Statistical analysis

To ensure statistical robustness, experiments were conducted for *n* = 1000 metacommunities, each of which had a unique spatial distribution of environmental variable and a unique set of randomly generated species, differing in environmental preferences and interspecies interactions. The assembly consisted of adding 600 species one by one to achieve a saturated metacommunity with the average alpha diversity of 29.99 and with gamma diversity of 53.29. Using this setup, we addressed the questions listed at the end of Introduction as follows:

i. Which credit metric most reliably quantifies progress towards an anticipated project outcome? To understand how predictably a given metric represents anticipated change in ecosystems resulting from a given intervention, we computed for each of the 12 metrics the coefficients of variation of metric change between the original and modified state resulting from each type of intervention, after subtracting results from the respective control conditions. The coefficient of variation represents the standardised measure of dispersion, calculated as the ratio of standard deviation to mean. It permits comparison of variability across metrics. For ease of comparison, the extinction ratio and STARt metrics were multiplied by –1 because for these two metrics larger values represent ecologically worse states. Project developers may favour the credit metric that exhibits the lowest coefficient of variation for the project’s type of restoration action.
ii. Which intervention most reliably generates gains in a given credit metric? This can also be addressed using the coefficients of variation computed above. Credit developers and investors confident in a credit metric’s utility, therefore aiming to design projects that reliably generate corresponding credits, may favour the type of restoration actions with the smallest coefficient of variation for the given credit metric.
iii. Are different aspects of biodiversity captured by different credit metrics? To investigate this question, pairwise Pearson’s correlations were computed of the changes between the original and modified states, after subtracting results from the respective control conditions, for all 12 metrics over the *n* = 1000 replicates, separately for each of the above-described nature perturbation experiment. The critical t-value at a 95% confidence level was calculated as t(0.975, df = 998) = 1.962. This corresponds to a critical threshold of correlation of approximately 0.062. This was followed by hierarchical cluster analysis to examine similarity between metrics. This analysis implements hierarchical clustering using the average-linkage to create a dendrogram which progressively merges observations based on their average dissimilarity until all elements are connected in a single tree structure (O’Sullivan et al., 2023). For ease of comparison, the extinction ratio and STARt metrics were multiplied by –1 because for these two metrics larger values represent ecologically worse states. To understand the metrics behaviour across and between the six nature perturbation experiments, hierarchical cluster analysis was also carried out on data pooled from all experiments, and on data pooled from all experiments with the mean response to the experiment subtracted.
iv. How well do certain credit metrics capture high-level objectives as quantified by global metrics such as the RLI, LPI or global MSA? This question is addressed using the pairwise correlation matrixes and cluster analyses computed above.

## 3. Results

Figure 1 shows the coefficients of variation (CV) of metrics, enabling comparison of variability across metrics, with each row representing an experiment and each column representing a metric. Lower coefficients indicate stronger correspondence between metric responses and perturbation effects, whereas values exceeding 1 represent unpredictability of metric responses (Fernández-Martínez et al., 2018). Experimental degradation is most closely tracked by biomass, RLI, LPI, MSA, and *credit metrics C* and *F*. While restoration experiments over the full or portions of model landscapes are tracked well by biomass, MSA, and *credit metrics C* and *F*, restoration of smaller project areas is best represented by biomass and *credit metric C*. Species-specific extinctions are most closely captured by the extinction ratio, LPI, MSA, and *credit metric C*. Species-specific restoration events are most closely tracked by LPI, while the environmental heterogeneity restoration experiments seem to only be tracked, if moderately, by c*redit metric F*. Metrics did not represent changes in only 1% of the project area well. However, this change already significantly when only 4% of the project area was altered.

**Figure 1.**
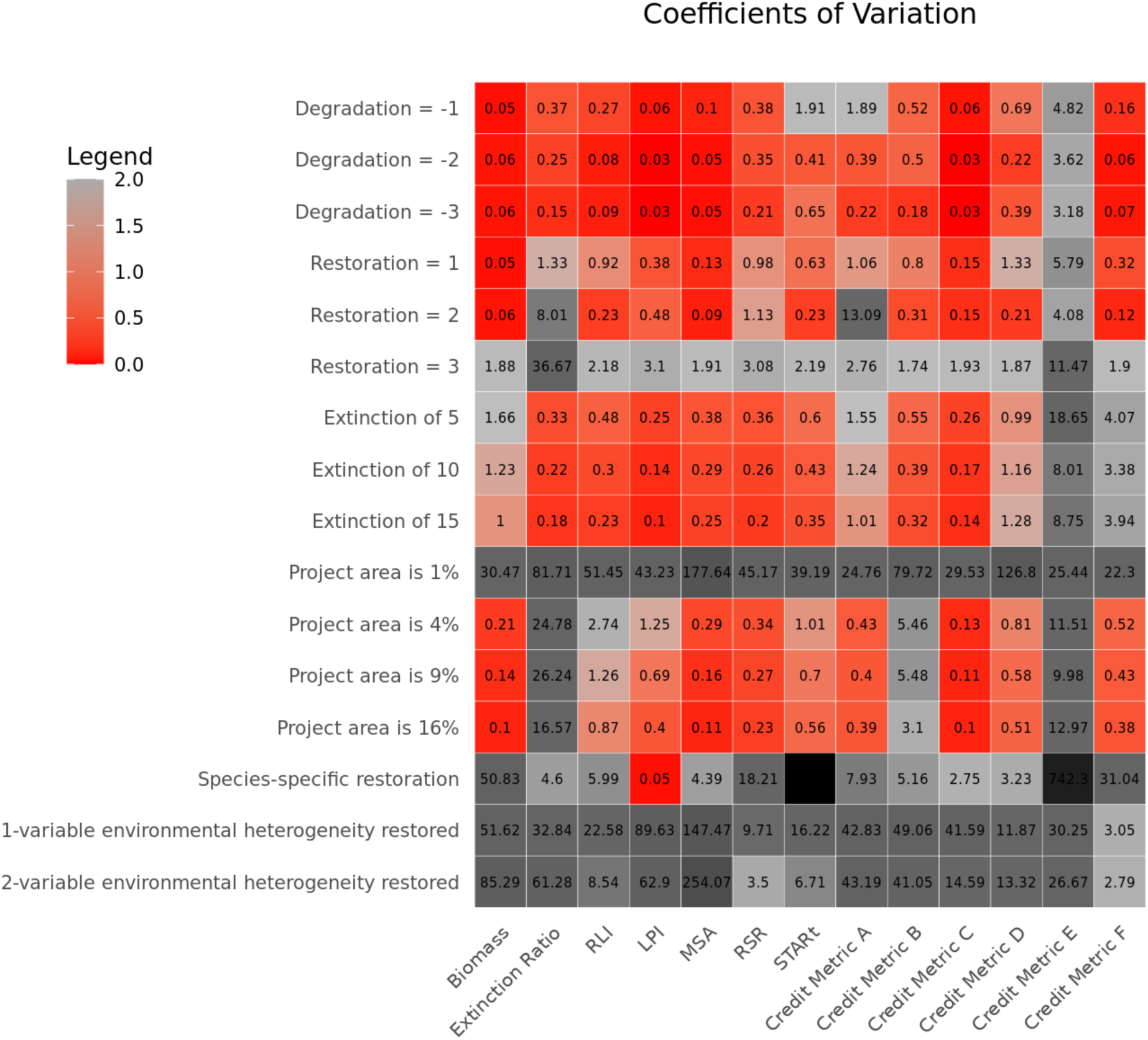
Coefficient of variation (CV) of metric responses across experimental settings for n=1000 replicates. The x-axis represents metrics; the y-axis represents experiments. CV values below 1 indicate stronger correspondence between metric responses and perturbation effects, while values above 1 suggest unpredictability. Due to high variability, the colour scale emphasizes informative CV values below 1. The legend spans only the small range, for larger values, refer to the numerical labels within the figure. To aid comparison, extinction ratio and STARt values were multiplied by –1, as higher values indicate ecologically worse states.

Figure 1 can also be used to determine which experiment is most aligned with a given metric, for instance, it suggest that *credit metric A* and *C* respond best to restoration of limited areas and *B, D* and *F* to broad restoration of moderately impacted habitat. LPI, on the other hand, responds most predictably to species-specific restoration..

Figure 2 shows the pairwise Pearson’s correlations of all 12 metrics’ n=1000 replicate experiments pooled over all nature perturbation experiments. This analysis emphasizes the between-experiment variation in the metrics’ behaviour, showing how the metrics behave compared to each other in response to different types of interventions. The observed patterns are dominated by the metrics’ mean responses to the perturbation experiments, especially where the CV of the responses is low. For this and all subsequent correlation analyses, we multiplied RLI and STARt with –1 as above. The results suggest that the credits’ behaviours fall into one of two clusters. One, aligned strongly with biomass change, contains MSA, LPI, RLI and *credit metrics C, D* and *F*. STARt also correlates with this cluster. The other cluster follows the extinction ratio, with strong correlations to RSR, MSA, RLI, and *credit metrics A and B*. Weaker correlations exist between these two groups, as the extinction ratio is weakly correlated with biomass. *Credit metric E* stands out by not falling into either cluster. *Credit metric C* is unique amongst credit metrics in having rather strong correlations with both biomass and extinction risk.

**Figure 2.**
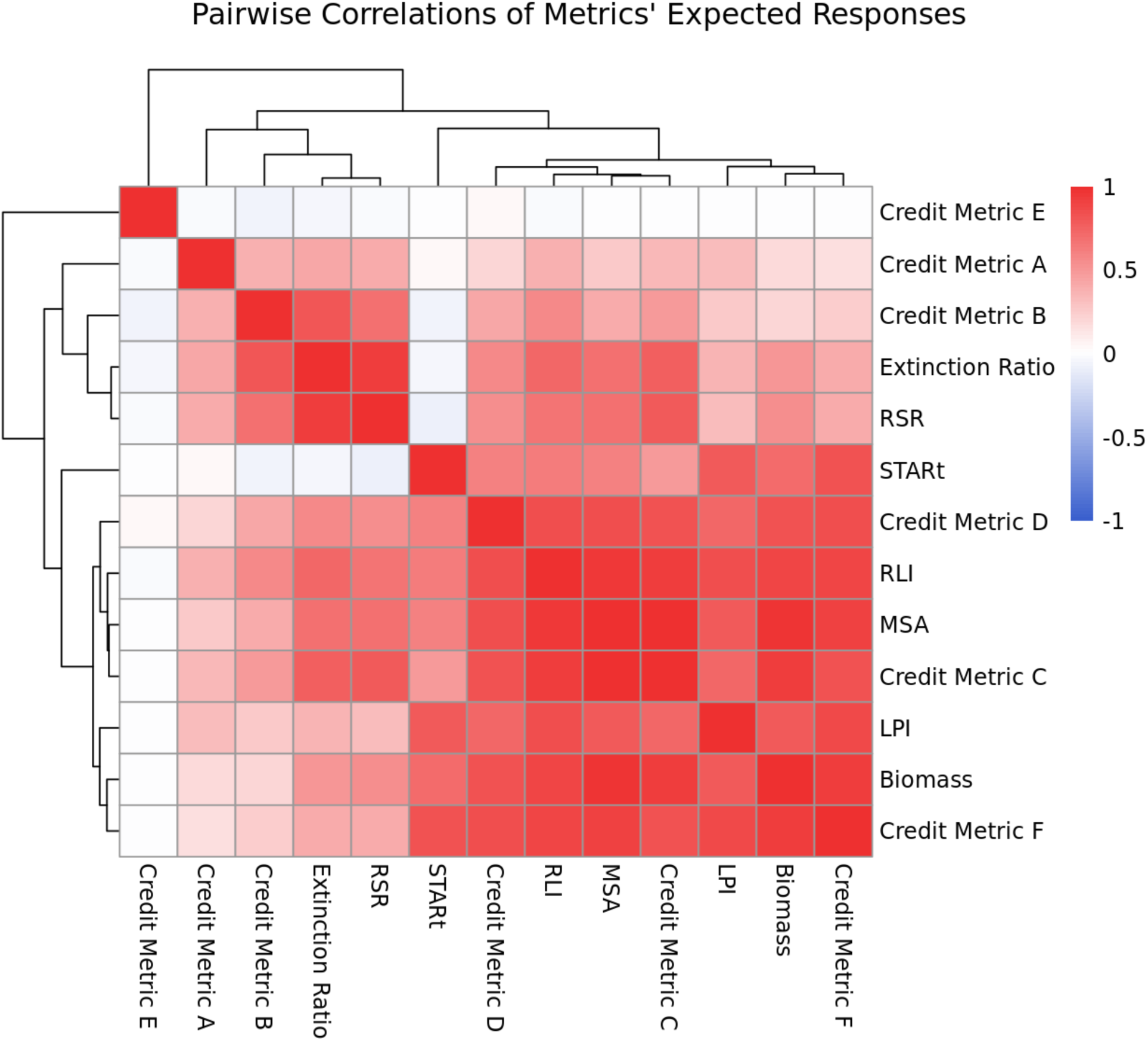
Pairwise Pearson correlations among 12 biodiversity metrics across 1,000 replicates pooled over all nature perturbation experiments, arranged via hierarchical clustering. To aid comparison, extinction ratio and STARt values were multiplied by –1, as higher values indicate ecologically worse states. Correlation patterns are primarily driven by metrics’ mean, expected responses to perturbations.

Figure 3 shows the result of an analysis analogous to that in Figure 2, with the only difference being that the mean response over the 1000 replicates is subtracted from the from the metric responses for each metric-experiment combination. This analysis summarises the within-experiment variations in metric behaviour over all experiments, probing for correlations in the metrics’ idiosyncratic responses to the random elements of experimental setups (distributions of environmental variables and the specific environmental preferences and interactions of species). Remarkably, the correlation structure seen in Figure 3 is very different from that in Figure 2. At least two distinct families of metrics can be distinguished. The first family comprises of two subclusters, one consisting of LPI, MSA, RLI and *credit metrics A* and *C*, the second of biomass, extinction ratio, RSR and *credit metric B*. The metrics across both these subclusters have higher sensitivity to decreases in abundances at the species-level. In detail: the extension ratio registers globally extinct species, LPI and MSA compare biomass declines to a predetermined base-state, RSR is a sum of all species’ presence-absence in a given area, *credit metric C* is a modification of MSA, *credit metric B* is the sum of the proportional changes in the global abundances of each species, and credit metric A is a product of area size and the sum of a wide variety of biodiversity metrics, including gamma biodiversity, Simpson’s Pielou’s and Whittaker indices. For ease of reference, we will refer to this cluster as *‘abundance-based’* metrics. Metrics in this family capture an element of extinction risk. The second family consists of *credit metrics E, F* and *D*. *Credit metrics E* and *F*, do not prescribe proxies for abundance of all species in their ‘basket of metrics’ methodologies. Instead, they and similarly *credit metric D*, recommend a selection of indicator species to synthesise an understanding of the ecosystem’s composition. For ease of reference, we will refer to this cluster as the *‘basket-of-indicator’* metrics.

**Figure 3.**
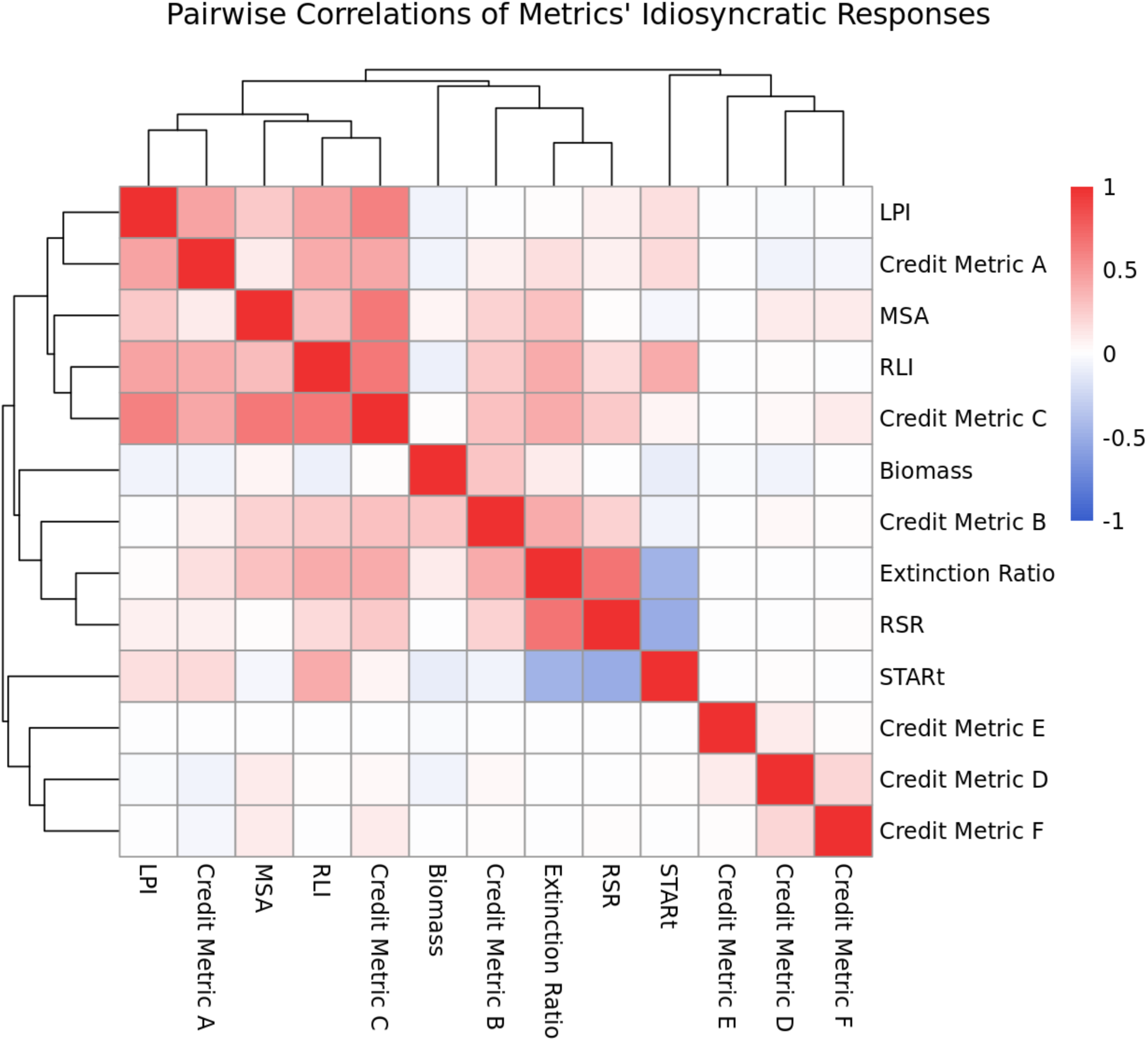
Pairwise Pearson correlations among 12 biodiversity metrics pooled across all nature perturbation experiments. Metric responses were mean-centred per metric– experiment combination across 1000 replicates to isolate idiosyncratic variation. Hierarchical clustering was used to arrange the correlation matrix. For comparability, extinction ratio and STARt values were multiplied by –1, as higher values indicate ecologically worse states.

Although RLI and STARt are both using IUCN Red List categories as the base of their methodology, their responses differ markedly in this analysis, with STARt displaying negative correlation with both the extinction ratio and RSR. The two metrics’ methodologies differ in several points. STARt is spatially explicit. When averaged over space and in absence of species extinctions, STARt divided by pristine gamma equals, up to linear transformations, RLI. However, while RLI incorporates historically extinct species STARt does not account for them because these are not associated with an extant populations’ location. This means that, if an extinction event occurs STARt scores will benefit, as a previously critically endangered species is no longer present. This effect reduces the correlation between STARt and RLI in Figure 3, which we’d otherwise expect to be strong. In situations where many extinctions occur, such as for the strongest level of environmental degradation we consider (Figure 4c), the effect can even lead to an anticorrelation between STARt and RLI. Similarity, as STARt is calculated for the project area, local removal of endangered species can reduce local STARt scores, even though lower STARt values are generally interpreted as preferrable over higher values. This explains STARt’s anticorrelation with the extinction ratio and RSR: every time an extinction or a local extirpation within the manged area occur STARt scores register an improvement.

**Figure 4.**
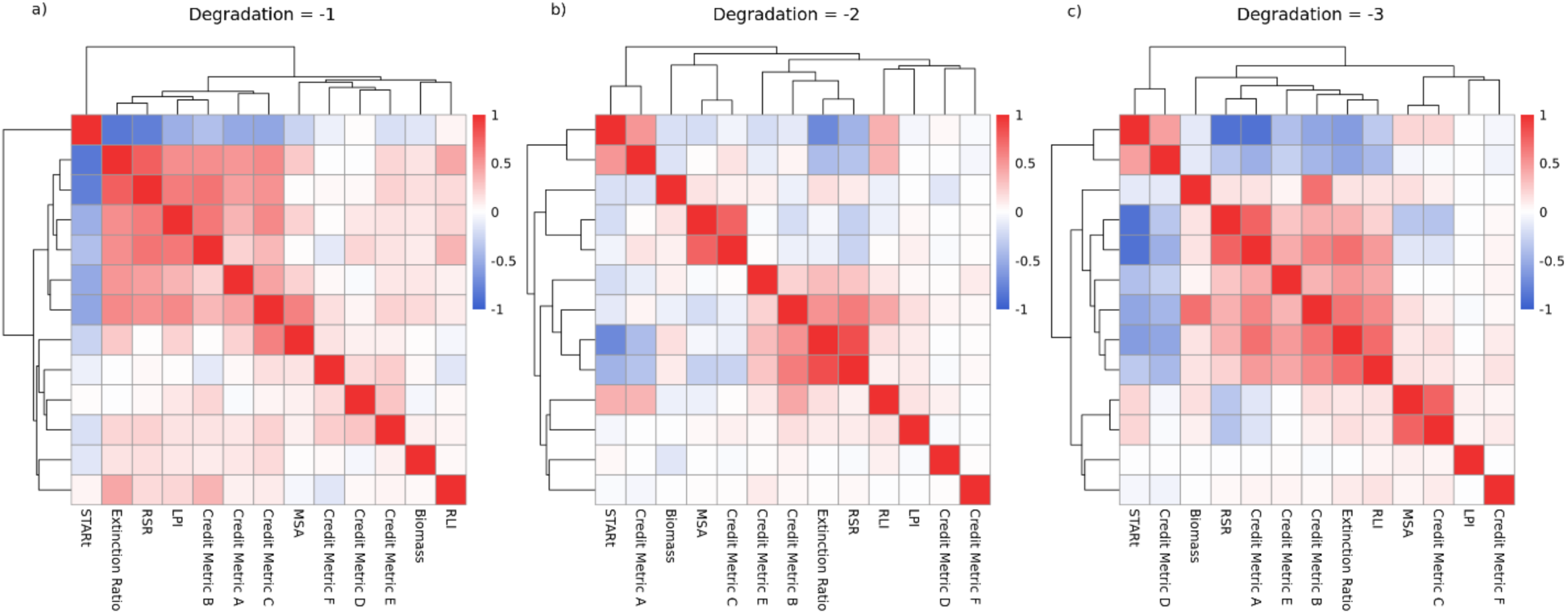
Pairwise Pearson correlations among 12 biodiversity metrics based on 1,000 responses to degradation experiments, arranged via hierarchical clustering. The three panels represent increasing levels of degradation. To aid comparison, extinction ratio and STARt values were multiplied by –1, as higher values indicate ecologically worse states.

Figures *4* to *7* display cluster analyses based on pairwise Pearson’s correlations of all 12 metrics’ n=1000 replicates, separately for each nature perturbation experiment. The above pattern separating the *‘abundance-based’* family of metrics from the ‘*basket-of-indicator’* family returns for many experiments. Specifically, the environmental restoration experiments (Figure 5), extinctions (Figure 6) restoration area increase (Figure 7a), and species-specific restoration (Figure 7b) reproduce this pattern, and it appears faintly in the environmental heterogeneity restored results (Figure 7c). In the case of the environmental degradation experiment (Figure 4) correlations seem to depend on the magnitude of impact, with the pattern more clearly reproduced for smaller perturbations. RLI and STARt display strong correlations when the entire simulated area is being manipulated and extinctions and local extinctions are limited, for instance, in the small magnitude environmental restoration (Figure 5a) experiment. Interestingly, STARt differs more markedly from RLI in the restoration area increase experiments (Figure 7a), where only a small percentage of the overall simulation area is being manipulated and evaluated by STARt. Here again, STARt displays negative correlation with metrics negatively accounting for extinctions, notably RSR, RLI and *credit metric B*.

**Figure 5.**
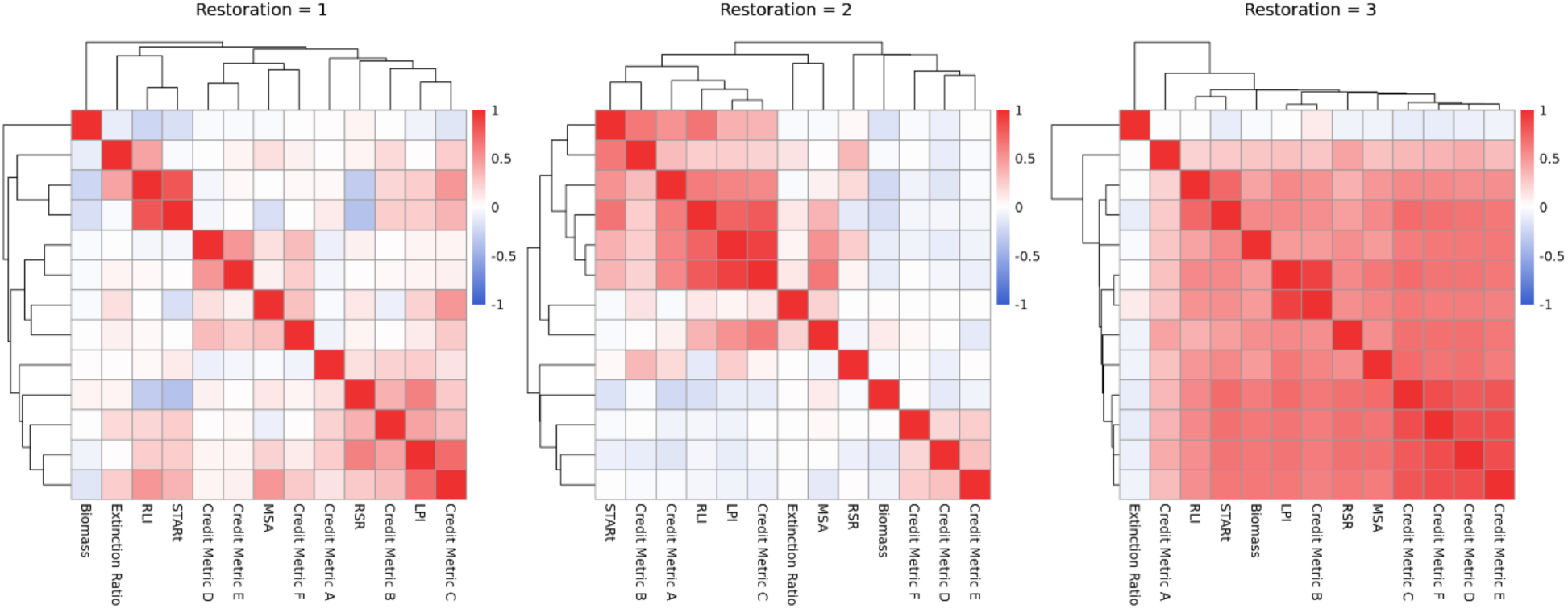
Pairwise Pearson correlations among 12 biodiversity metrics based on 1,000 responses to restoration experiments, arranged via hierarchical clustering. The three panels represent increasing levels of degradation being restored. To aid comparison, extinction ratio and STARt values were multiplied by –1, as higher values indicate ecologically worse states.

**Figure 6.**
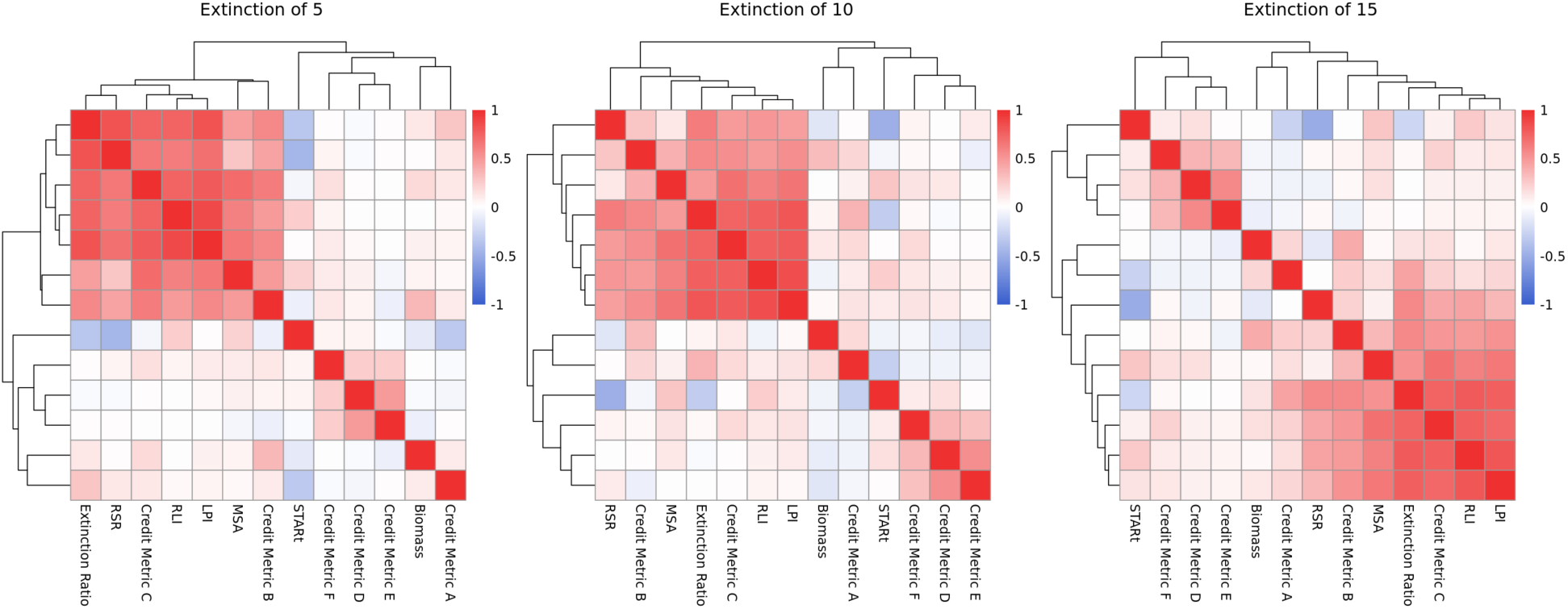
Pairwise Pearson correlations among 12 biodiversity metrics based on 1,000 responses to species-specific extinction experiments, arranged via hierarchical clustering. The three panels represent increasing numbers of extinctions. To aid comparison, extinction ratio and STARt values were multiplied by –1, as higher values indicate ecologically worse states.

**Figure 7.**
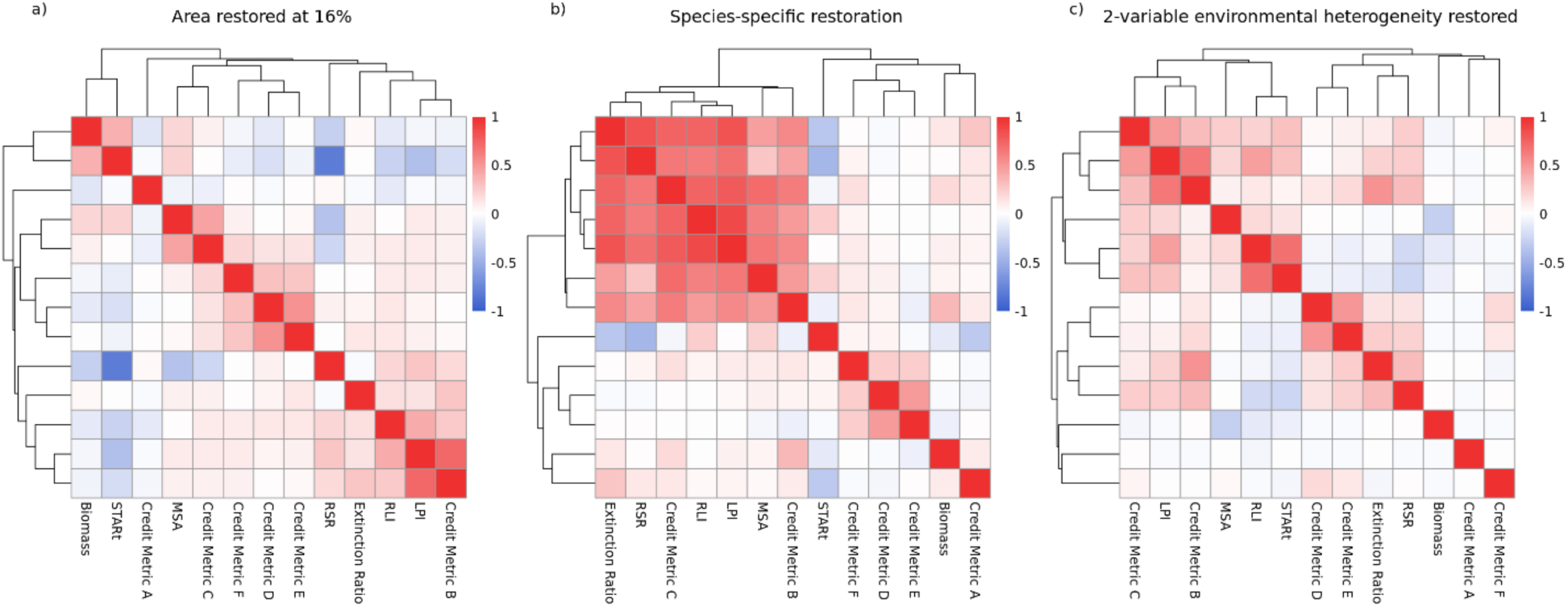
Pairwise Pearson correlations among 12 biodiversity metrics based on 1,000 responses to three distinct nature perturbation experiments, arranged via hierarchical clustering. (a) Simulated restoration of 16% of global area. (b) Restoration targeting a single critically endangered species. (c) Restoration of environmental heterogeneity modelled with two variables. To aid comparison, extinction ratio and STARt values were multiplied by –1, as higher values indicate ecologically worse states.

**Figure 8.**
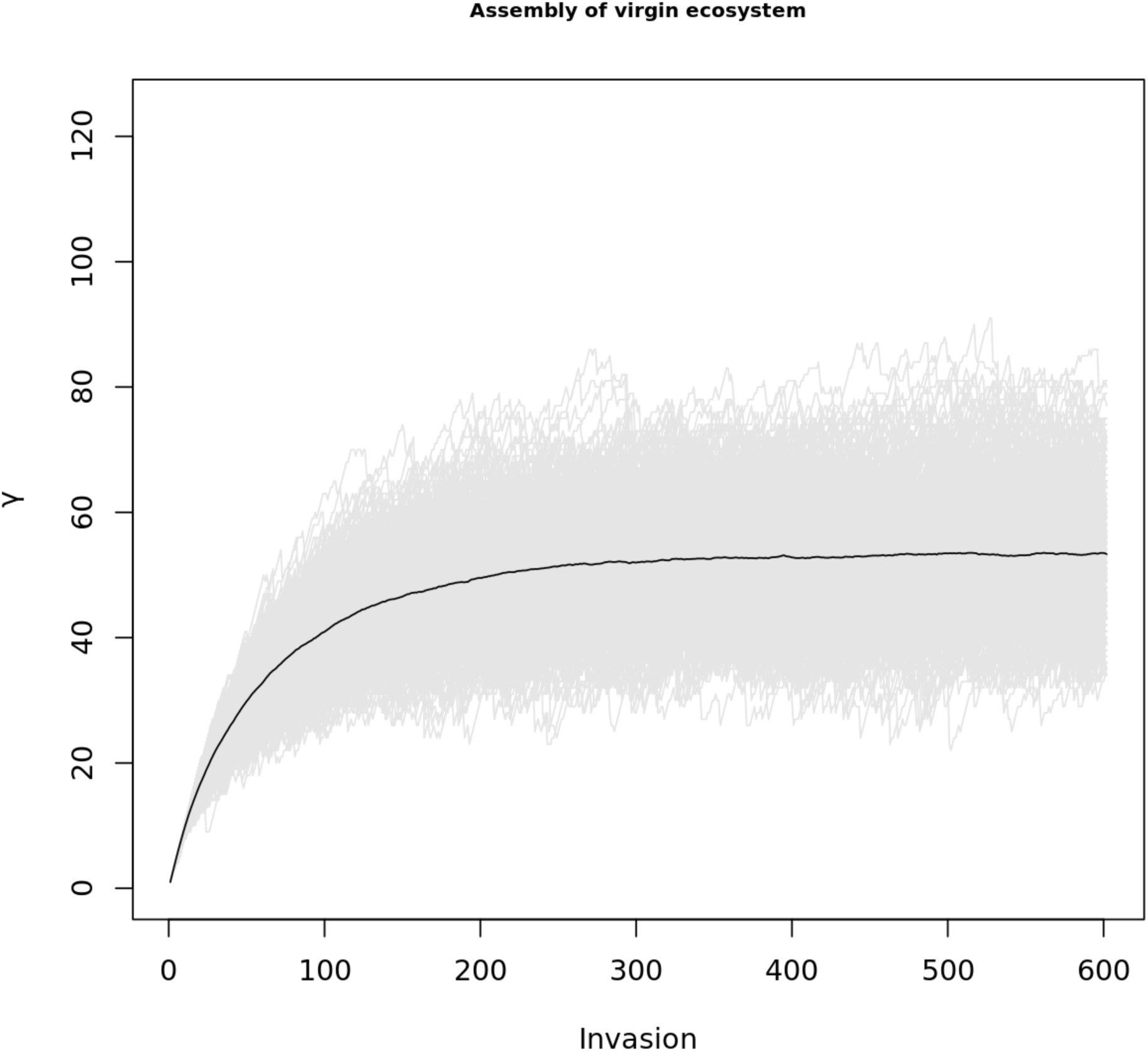
illustrates the saturation of regional species richness. The vertical axis represents gamma diversity, while the horizontal axis corresponds to the cumulative number of invasion events. Saturated metacommunities generated through this process were found to replicate a range of landscape-level ecological patterns observed in empirical systems (O’Sullivan et al., 2019, 2021, 2023). Accordingly, these saturated states are employed in this study to represent undisturbed, pristine ecological communities.

These results suggest that the metrics’ responses are heavily dependent on the type of nature perturbation carried out. For instance, credit metric A shows positive correlations with biomass in species-specific extinction, restoration area increase, and species-specific restoration experiments but no correlation in environmental restoration or degradation experiments. Similarly, while MSA and *credit metric C* are highly correlated for most experiments (Figures 4, 5, 6, 7a, 7b), they are not in the environmental heterogeneity restoration (Figure 7c). Furthermore, in the environmental degradation experiment (Figure 4) results are markedly different depending on the degree of degradation simulated. While Figure 4a has the two distinct clusters, these do not arise for more severe environmental degradation. By contrast, species-specific extinction experiments (Figure 6) lead to similar grouping patterns with increasing intensity, i.e. increasing numbers of extinctions. LPI, MSA, RLI, RSR and *credit metric C* form one of these groupings, while *credit metrics D*, *E* and *F* form another.

In Figure 4c the severity of the degradation has led to the catastrophic collapse of the ecosystem, which is attempted to be restored in Figure 5c; therefore, the results from these conditions might not be applicable to normal circumstances. In the restoration area increase (Figure 7a), species-specific restoration (Figure 7b) and the environmental heterogeneity restoration (Figure 7c) experiments the variation of the respective experimental conditions (area size, magnitude of perturbation, and number of environmental variables) did not impact the correlation clusters, only the correlation strength. Therefore, only the experimental conditions with the strongest correlations are shown in Figure 7 (area at 16% of global; species-specific restoration = 0.05; 2-variable environmental heterogeneity).

## 4. Discussion

### 4.1. Interpretations of key findings

This paper investigated how twelve metrics proposed for the quantification of biodiversity, six of which form the basis of biodiversity crediting schemes, responded to a set of nature perturbation experiments representing human interventions into nature. We aimed to shed light on not only the modelled metrics’ characteristics, but also onto what components of biodiversity influences their behaviour. To do so, we outlined four fundamental questions. As answers, our results suggest that (1) metrics respond differently to diverse forms of ecosystem perturbations, therefore they clearly must capture different elements of biodiversity or emphasise these elements differently. (2) These differences translate to meaningful differences in metric behaviour, as certain metrics are better suited to quantify the effects of certain interventions, and vice versa; certain types of interventions are better suited to generate certain metrics (Figure 1).

(3) Quantitative comparability of credits based on different metrics is limited. While the metrics modelled cluster into correlated categories, one needs to distinguish between correlations in how metrics respond to various specific manipulations or project types and correlations in idiosyncratic metric dependencies on system details. Correlations in the predictable, expected aspects of a metric’s response to a given manipulation are illustrated in Figure 2, while Figure 3 characterises the idiosyncratic aspects. Metrics that correlated in

Figure 2 are expected to respond to different nature manipulations in a similar fashion. Therefore, Figure 2, in conjunction with the CVs in Figure 1, may support project developers at the stage of deciding which metric to use and what kind of project to develop. As there are two clusters here, one correlated with biomass and another with extinction ratio, our results suggest that project developers may consider a selection of metrics capturing both biomass and extinction risk for their projects.

Conversely, metrics that are correlated in Figure 3, thus capturing the idiosyncratic behaviour of the system in a similar fashion, respond similarly to variations in detailed ecosystem properties. To the extent that these properties are controllable by the management of ecosystems, Figure 3 may therefore help users understand the ecosystem management style that a given metric incentivises. By our understanding, incentivising particular management styles by project developers is one of the intended benefits of assigning biodiversity credits based on measured rather than expected ecological outcomes, which is why the alignment of metrics shown in Figure 3 is just as important for understanding biodiversity credit as the somewhat different alignment of expected responses underlying Figure 2. Based on their idiosyncratic responses, the metrics in our study fall into two families: ‘*abundance-based’* metrics, which have a higher sensitivity to changes in abundance at the species level; and ‘*basket-of-indicator’* metrics, which describe the ecosystem through a selection of indicator species. Some credit metrics were intentionally designed to align with established indicators, accounting for their placement in the above families. Credit metric C is a project-specific adaptation of MSA, and metric B similarly modifies LPI, accounting for their frequent close correlations. STARt was developed to approximate RLI, which can lead to strong correlations in the absence of extinctions or extirpations. Metrics D, E and F follow basket-of-indicator methodologies, explaining their, albeit modest, positive correlations.

The differences in correlation patterns between Figures 2 and 3 reflect the sensitivity of metrics to the details of their definitions. For instance, average responses of STARt align with changes in biomass but shows no correlation with extinction ratio (Figure 2). However, STARt’s response to idiosyncratic variation between ecosystems displays a rather strong anticorrelation with the extinction ratio. This likely reflects that STARt values can decrease due to both an improvement in a threatened species’ IUCN Red List categorisation, or the removal or extinction of a threatened species from the area of consideration.

Finally, we endeavoured to answer (4) whether generation of credits directly contributes to globally agreed objectives, by examining whether credit metrics are correlated with metrics designed to quantify high-level objectives (e.g., LPI, MSA, RLI). As demonstrated above; to be truly aligned with these metrics, credit metrics need to show correlations in both in the expected and idiosyncratic aspects of their responses, which may differ significantly between metrics (Figures 2 and 3). Nearly all credit metrics’ expected responses correlated with a metric designed to quantify high-level globally agreed objectives. However, only metrics in the *‘abundance-based’* cluster had the idiosyncratic aspects of their responses correlate with metrics designed to quantify high-level globally agreed objectives.

Nevertheless, as Figure 1 indicates, metrics from all clusters may be able to closely track environmental degradation and restoration. This relationship may be attributable to a modest positive correlation between extinction risk and biomass (Figure 2); as if biomass increases across species, the extinction ratio tends to decline. Consequently, metrics that quantify extinction risk may also serve as indirect indicators of ecosystem status.

Biodiversity is highly complex and multidimensional and may be conceptualised in a wide variety of manners based on socio-cultural contexts and considerations (Pascual et al., 2021; Soto-Navarro et al., 2021). Projects may focus on differing objectives such as maximising ecosystem integrity, species conservation, or ecosystem services. Accordingly, credit designers need to make the choices to consider a limited number of such aspects of biodiversity, especially in a bid to remain simple to comprehend and be adopted by project developers. All biodiversity metrics must effectively translate intricate ecological information into accessible quantitative scales to avoid the systematic undervaluation of nature in regulatory and corporate decision-making (Strange et al., 2024). Inherently, these differing decisions prime metrics to have varying sensitivities when subjected to a wide enough range of ecosystem perturbations.

Indeed, we have found that most metrics are either attempting to capture the ecosystem’s status through a measure of biomass or are estimating the extent of extinction risk at the species level. Moreover, our results suggest that these two clusters exhibit different responses to a range of ecological interventions. These different approaches represent distinct methodological paradigms which incentivise two very different ways of choosing, designing, and managing projects. Any attempts to use both may require trade-offs. This suggests that a singular metric may not have the sufficient scope to comprehensively capture both an estimation of ecosystem service through biomass and species-level extinction risk trajectories simultaneously. We suspect that the fundamental divergence arises from competing temporal priorities: maximising immediate ecosystem service extraction versus ensuring indefinite species preservation (Everitt, 1980). Previous research suggests that maximising short-term ecosystem services often may be at odds with the goals of long-term species conservation (Clémençon, 2021; Qiu & Mitchell, 2024). This dichotomy necessitates specialised metric development for either approach, rendering the prospect of a universally applicable biodiversity credit metric improbable, and highlighting the equal importance of metrics estimating ecosystem state and extinction risk.

### 4.2. Limitations

The coefficient of variation (CV) analysis indicates how consistently a metric responds to changes in the experimental variable. However, CV generally depends on the magnitude of the intervention, which limits its comparability across different intervention types. When the expected response of a metric is zero or near zero, CV can become disproportionately large within this range but smaller at other intervention magnitudes. This variability further constrains the interpretability and comparability of CV in this study. Specifically in the species-specific restoration experiment, a minor change to any one species should benefit the entire community; however, above a certain threshold, the success of the targeted species may adversely affect other species, resulting in an overall detrimental impact on the community. The coefficient of variation analysis is not able to pick up this nuance, therefore suggesting that *credit metric B* is not suitable for the experiment, despite it reproducing the expected pattern.

As described in Methods, the global area of the simulations was relatively small when compared to the mean range of species which successfully got established in the assembled communities. Therefore, our results overestimate the risks of primary and secondary extinctions resulting from our experimental interventions. This is counterbalanced by the fact that the modelled abiotic environment is not overly complex, supporting an average alpha diversity of 29.99 and gamma diversity of 53.29. To model the effects of environmental degradation and restoration the species’ growth rates were modified directly rather than the environmental variables. This means that in the degradation and restoration phases of an experiment all species were equally impacted. Therefore, a species which had strongly favourable conditions in the undisturbed environment, and therefore had a high growth rate, could still retain a positive growth rate in the event of a minor decrease of all species’ growth rates. This methodology is representative in case of small deviations, however, would lead to a full collapse of the community in extreme degradation cases. In nature, such extreme degradation cases would not always lead to the complete collapse of the ecosystem. Rather, depending on the nature of the degradation, it may lead to the emergence of very few generalists which are able to cope with the degraded environment.

### 4.3. Opportunities for further research

Future modelling efforts could incorporate credit metrics based on ecosystem connectivity, habitat fragmentation, or taxonomic dissimilarity. Additionally, we recommend conducting large-scale simulation studies on landscapes that are substantially larger than species’ ranges and where average species ranges are large relative to project area sizes.

Experimental and modelling studies could investigate metrics responses to offsetting experiments. We hypothesise that metrics estimating species extinction risk or biomass might have differing responses. A literature review could shed further light on the role of ecosystem services and extinction risk in biodiversity metrics, and the nature markets. Finally, studies should investigate how government action can incentivise the acceleration private finance in nature.

### 4.4. Implications

Biodiversity credits are financial instruments that represent evidence-based units of positive biodiversity outcomes. In the carbon markets, credits are standardised and fungible, with each credit corresponding to one metric tonne of CO₂ removed from the atmosphere. This fungibility underpins buyers’ claims of net-zero emissions. For biodiversity, establishing similar fungibility is more complex, as it depends on demonstrating equivalence between ecological degradation at one site and restoration at another(Bezombes et al., 2017; zu Ermgassen et al., 2020; zu Ermgassen & Löfqvist, 2024). To deal with this issue, applications of biodiversity credits vary heavily. Use-cases may range from merely measuring the impacts of voluntary contributions to nature, voluntary compensations of impacts throughout a business’ value chain, or attempting to offset the specific impacts of development projects mandated by regulation. Credit schemes therefore differ in the project activities they offer, which can range from the measured and evidenced maintenance of high value biodiversity areas through measures of ecosystem services, to active restoration of desirable habitats and ecosystems through measures of extinction risk. In this context, our results suggest significant differences in the way metrics respond to interventions and imply that a universal biodiversity credit is improbable. Other studies have questioned the benefits of establishing a universal biodiversity credit, when accounting for engrained uncertainties in ecosystem status metrics (Wauchope et al., 2024). Therefore, projects should use metrics which are specifically suitable for their ecological aims, compatible with their project types and evidencable for their financial use-cases.

Our results further illustrate this point through the simulated metrics’ limited ability to detect changes as a response to the restoration of environmental heterogeneity experiments. This might mean that no change has occurred as a result in the ecosystems, or that the metrics simulated here are missing the changing aspect of biodiversity. We suspect the latter, as none of the simulated metrics include a spatial component, such as connectivity or habitat fragmentation. Such credit metrics exist, for instance *Plan Vivo’s Biodiversity Certificates* or *Terrasos’s Biodiversity Credits*. Unfortunately, their inclusion in this study is currently limited by the complexity of implementation, due to both the computation of spatial component and other aspects such as calculating taxonomic dissimilarity. Nonetheless, this result highlights the vulnerabilities of using an ill-suited credit metric to measure the impacts of interventions in nature.

Despite the studied credit metrics’ observed differences in methodology and behaviour, outcome comparability across projects is essential for a functional biodiversity credit market. Credit buyers must be able to evaluate and contrast schemes to identify those aligned with their objectives. Our findings indicate that comparability should be pursued among projects targeting similar ecological outcomes. Specifically, those aiming to reduce extinction risk should be assessed relative to others with the same goal, while projects enhancing ecosystem services should be compared within that category.

Furthermore, our results also suggest that to ensure alignment with Goal A and Target 4 of the Kunming-Montreal Global Biodiversity Framework on halting species extinctions, projects should also consider a metric sensitive to decreases in species-level abundances and thereby reflecting extinction risk. This is crucial, as metrics based on the number of extinctions only capture past, irreversible losses. In contrast, metrics that can capture elevated extinction risk offer early warnings, enabling proactive conservation before species loss occurs.

Our findings highlight key considerations for the development of biodiversity credit markets. If extinction risk and ecosystem service metrics reflect different elements of a successful restoration project, credit suppliers must determine how to best bring these elements to market. For projects targeting a single outcome (e.g. increasing ecosystem services), perhaps a single corresponding metric may suffice. However, when multiple outcomes are achieved, such as simultaneous gains in ecosystem services and reductions in extinction risk, it remains unclear whether these should be bundled into a unified credit or offered separately as a stack (Robertson et al., 2014). While such questions are central to the long-term design of biodiversity markets, they introduce additional complexity. Early-stage projects and transactions may benefit from prioritising simplicity and ensuring transparency regarding what each credit represents.

## 5. Supporting Information

### 5.1. Lotka – Volterra Metacommunity Model (LVMCM)

The configuration of the LVMCM followed the protocol outlined by O’Sullivan et al. (2023) and all model parameters are as in O’Sullivan et al. (2023) if not otherwise stated. A detailed description of the model components is provided below.

#### 5.1.1. Environmental heterogeneity and spatial structure

Environmental heterogeneity was represented by assigning to each site *x* a set of *K* independent random variables E_kx_ *(1 ≤ k ≤ K).* These correspond to principal components of abiotic environmental variation. These variables were drawn from spatially correlated two-dimensional Gaussian random fields, characterised by a mean of 0, a standard deviation of 1, and a correlation length of 1.

Each species *i* was assigned an environmental optimum 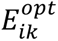 for each environmental variable *k*, sampled from a uniform distribution over the interval: 1.5 * min(*x*) E_kx_; 1.25 * max(*x*) E_kx_. The degree of environmental heterogeneity experienced by species was modulated by the niche width parameter *w*. The intrinsic growth rate of species *i* at site *x* was calculated using the following expression:

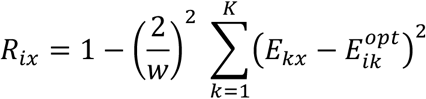

To simplify the parameterisation, all species were assumed to share identical niche widths across all *n* independent environmental components. In the metacommunity simulations presented in this study K was set to either 1 or 2. The spatial network structure of sites varied across experimental setups and is described in detail within the respective methods sections.

#### 5.1.2. Metacommunity dynamics

The dynamics of the metacommunity were simulated using a spatially explicit extension of the classical Lotka–Volterra community model, following previous works of O’Sullivan et al. (2019, 2021). The formula is the following:

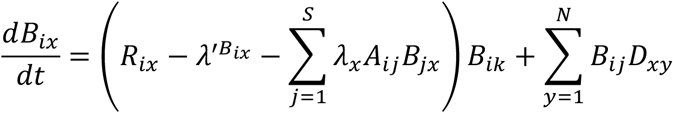

In this formulation, *B_ix_* denotes the biomass of species *i* at site *x*, while *R_ix_* represents the intrinsic growth rate of species *i* at site *x*, which varies spatially across the landscape.

To simplify the interaction structure, the unscaled local interspecific interaction coefficients *A_ij_* were drawn from a discrete probability distribution defined by *P(A_ij_=0.3)=0.3*, with *A_ij_=0* otherwise.

Spatial dispersal between sites was governed by the connectivity matrix *D_xy_*, where emigration and immigration rates were specified as follows: for each site *x*, *D_xx_= − e*; for connected sites *x* and *y*, 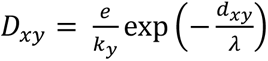; and D _xy_=0 for all other site pairs. Here, *e* denotes the emigration rate, d_xy_ is the Euclidean distance between sites *x* and *y*, and λ is the characteristic dispersal length, which was held constant throughout this thesis. The term *k_y_* refers to the unweighted degree of site *y*, serving as a normalisation factor.

#### 5.1.3. Metacommunity assembly

Model metacommunities were constructed through an iterative invasion process. At each step, a single new species *i=1* was generated with randomly sampled environmental preferences and interaction coefficients *A_ij_*, and introduced into the system at low biomass. The community dynamics were then simulated over a period of 3000 unit times using the governing equation described above. Prior to each invasion event, the metacommunity was examined for species whose biomass had declined below the extinction threshold of 10^−4^ units across all sites. Such species were deemed regionally extinct and subsequently removed from the system.

This assembly procedure led to saturation of the metacommunity in terms of both local (alpha) and regional (gamma) species richness (O’Sullivan et al., 2019), a phenomenon attributed to the emergence of ecological structural instability (see Figure 8) (Rossberg, 2013). Once saturation was reached, each subsequent invasion resulted in the extinction of one resident species on average.

### 5.2. Biodiversity credit implementation

#### 5.2.1. BioCarbon Registry’s Biodiversity Credit

BioCarbon Registry conceptualise their Biodiversity Credit as a unit of measurement quantifying biodiversity gains at a given area. The number of credits issued is a product of area size and the sum of a wide range of categorised biodiversity metrics. These metrics are the following: Landscape Biodiversity Index, Species richness, Magalef Index of Diversity, Simpson’s diversity index, Pielou’s Index, Jaccard similarity index, Whittaker Index, Gamma index, High Conservation Values, Number of species in different IUCN Red List categories. For all the above-mentioned metrics BioCarbon Registry’s Methodology Paper provides a set of threshold values which assign a numerical factor, to be used in the sum calculation. After conducting conservation activities and assessments, Biodiversity Credits are issued in response to the positive change in metrics in the project area.

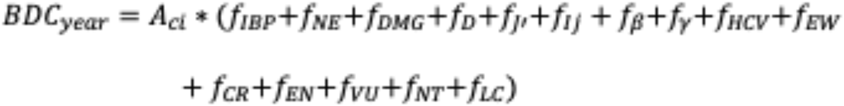

Where:

*BDC*_year_ = Biodiversity credits for a given amount of years

*A*_ci_ = The area within the boundaries of the conservation initiative; in hectares

*f*_IBP_ = Landscape Biodiversity Index

*f*_NE_ = Specific richness (number of species)

*f*_DMG_ = Margalef Index of Diversity

*f*_D_ = Dominance index (or Simpson’s diversity index)

*f*_J’_ = Pileou’s Index

*f*_Ij_ = Jaccard similarity index

*f*_β_ = Whittaker index

*f*_y_ = Gamma index

*f*_HCV_ = High Conservation Values

*f*_EW_ = IUCN Red List’s Extinct in Wild

*f*_CR_ = IUCN Red List’s Critically Endangered

*f*_EN_ = IUCN Red List’s Endangered

*f*_VU_ = IUCN Red List’s Vulnerable

*f*_NT_ = IUCN Red List’s Near Threatened

*f*_LC_ = IUCN Red List’s Least Concern

Implementation in the LVMCM:

The majority of the biodiversity metrics required by the BioCarbon Registry Biodiversity Credit are tracked straightforwardly by the LVMCM, these are: Species richness, Magalef Index of Diversity, Simpson’s diversity index, Pielou’s Index, Whittaker Index, Gamma index, Number of species in different IUCN Red List categories. Further information on how species are assigned IUCN Red List Categories can be found below. Some of the High Conservation Values (HCVs) must be assessed on a project-by-project basis or are not possible to model in the LVMCM, for instance community needs or cultural values. Other conditions are almost always guaranteed within the model, for instance areas of high species diversity. Therefore, the score for HCVs is guaranteed to be between one and three in the LVMCM, and accordingly the factor allocated is 1.02 and is kept constant, as per the methodology paper.

In BioCarbon Registry’s Biodiversity Credit methodology, the Jaccard similarity value is used to evaluate the similarity between multiple project locations. The LVMCM simulates a continuous project area without segmentation, therefore the Jaccard similarity is kept constant at 0.1, and therefore the factor allocated is 1.05.

The Landscape Biodiversity Factor cannot be calculated in the model, as landscape and area configurations are manually provided from outside the model. Therefore, the factor is kept constant at 1.

After these minor modifications the issuable Biodiversity Credits are calculated as the product of area size and the change in the sum of all biodiversity metrics.

#### 5.2.2. Botanic Gardens Conservation International’s Biodiversity Impact Credits

The Biodiversity Impact Credits (BICs) quantify the changes in the global species extinction risk caused by human interventions to the natural environment. Calculating BICs require the current global population size, the change in the local population affected by the intervention and a regularisation constant representing the risk of random extinctions at low population sizes. The BIC for a single species is the change in regional abundance, divided by the sum of the global population and a regularisation constant. Often the regularisation constant is negligibly small compared to the global abundance. BICs for a project area is the sum of all species-specific BICs.

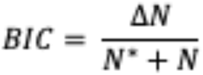

Where:

*N* = the current global population size of the species

*ΔN* = the change in the population of the species that resulted from the intervention

*N*∗ = regularisation constant as small population sizes a species can go extinct if by chance

Implementation in the LVMCM:

As the LVMCM tracks the exact abundances for all species both in and outside of the project area calculating the BICs is straightforward. The regularisation constant was chosen at 1% of the monoculture carrying capacity of a single site and kept constant.

#### 5.2.3. Organisation for Biodiversity Certificates’ Biodiversity Index

Organisation for Biodiversity Certificate’s Biodiversity Index Assessment Method describes a modified mean species abundance (MSA) calculation methodology. The carrying capacity of the project area is estimated by external experts at each assessment, against a predefined reference condition. In the case of the unmodified MSA calculations this reference condition is the ‘undisturbed’ state of an ecosystem. Assigning such an undisturbed state in practice can be challenging. Moreover, if the plan of restoration is different from the undisturbed state of the ecosystem, the assigned reference values might not even capture the full extent of the restoration efforts. The Organisation for Biodiversity Certificate’s Biodiversity Index (BI) circumvents these potential issues by allowing for expert aided free assignment of the BI reference value on a scale of zero to one. Subsequently, the BI is measured at each assessment, and credits can be issued as a product of the additional BI gained compared to the baseline measurement, and project area size.

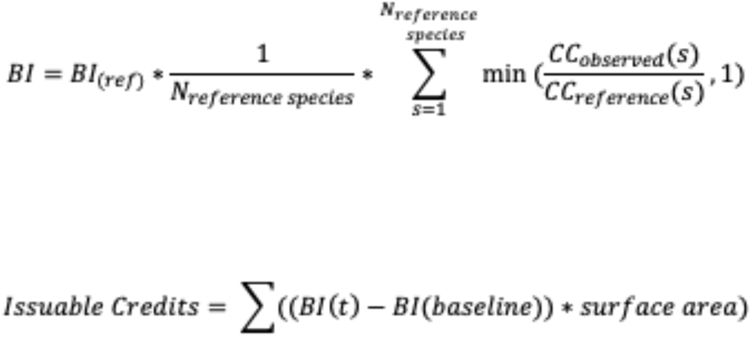

Where:

*BI* = Biodiversity Index

*BI*_(ref)_ = Biodiversity Index of the reference ecosystem

*N*_reference species_ = Total number of species in the reference ecosystem

*CC*_observed_(s) = Carrying capacity of species s in the assessed ecosystem

*CC*_reference_(s) = Carrying capacity of species s in the reference ecosystem

Implementation in the LVMCM:

For the purposes of the model, carrying capacity has been defined as biomass achieved by species s at a given location. As the LVMCM grants the opportunity of observing the same site in different conditions, the reference site is defined as the project site in its undisturbed state immediately after assembly. The BI reference value has been set at one and kept constant, to allow for an unconfounded comparison with alternative credits. Subsequently, the BI values were computed for the degraded and restored states, and the credits are calculated as the product of the difference between the degraded and restored states, and project area size.

#### 5.2.4. Savimbo’s Biodiversity Credit

Savimbo’s Biodiversity Credit (VBC) is a sum of the difference in integrity over time and area.

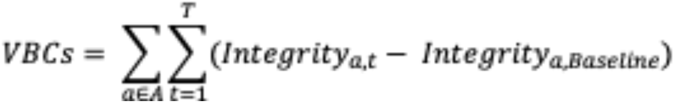

Where:

*VBCs* = Voluntary Biodiversity Credits

*a* = project area

*t* = time passed

Integrity is defined on a spectrum between 0 and 1 where 0 is a completely degraded ecosystem while 1 is a fully intact ecosystem. Integrity is sampled by the presence or absence of indicator species which are chosen by the user to represent the relative health of the ecosystem. If a species is representative a fully intact ecosystem it is assigned 1, however species which can survive in more degraded environments are only assigned a fractional “Integrity” score. These are summed for each measurement point within the project area; however, the sums cannot exceed 1. Change in “Integrity” is then calculated by subtracting the baseline “Integrity” figure from the changed one.

Furthermore, Savimbo track the “Value” of an ecosystem, which influences the quality of their credits issued. Ecosystem value is determined by public ecosystem indicators. These can be local indicator databases or the IUCN Red List of Ecosystems database.

Implementation in the LVMCM:

In the model, the integrity calculations rely on indicator species. Their selection in the model is described below. To compute a site’s integrity score within the project area, the model checks whether any of the three indicator species are present and awards a 0.5 integrity for each indicator species for the site. These values were chosen following Savimbo’s publicly available examples of integrity scoring. These are summed and truncated at 1 for a given site, in accordance with the Savimbo methodology. The site-wise Integrity scores are summed over the project area to calculate the Integrity for a given state in time allowing for the estimation of “Integrity”.

For the Ecosystem value term, the IUCN Red List of Ecosystems database is modelled for each assembled community. Further information on how the IUCN Red List of Ecosystems categories are assigned can be found below in section 2.4.2.

#### 5.2.5. Verra’s Nature Credit

The Nature Credit is defined as the biodiversity uplift of one “quality hectare equivalent” relative to a baseline, due to a project intervention. Quality hectares are a weighted unit of the area size and condition. At the project start, and at each following assessment, the condition of an area is estimated by the arithmetic mean of at least 3 structure and at least 2 composition indicators. In Verra’s methodology structure indicators are defined as “biotic or physical size and form, physical and chemical characteristics” with examples such as total biomass, canopy cover, water chemistry. While composition indicators are defined as the “variety, identity, and abundance of organisms” with examples such as species richness of characteristic biota or abundance of keystone species. All condition indicators are standardised against a reference site. Subsequently, the net biodiversity impact is calculated by comparing the condition-adjusted area figures at the project start with those at the current time, adjusting for leakage and the area’s global significance. After adjusting by an area-specific buffer value, the Nature Credits are ready to issue.

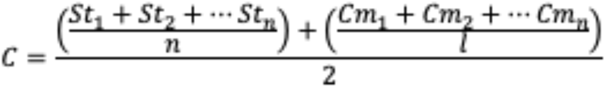

Where:

*C* = Condition indicator at a given time

*St* = Standardised structure indicators at the given time

*Cm* = Standardised composition indicators the given time

*n* = Number of structure indicators

*l* = Number of composition indicators

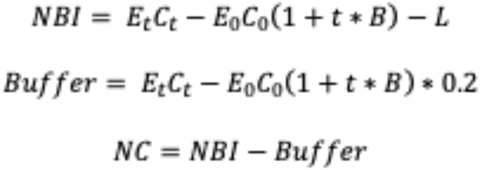

Where:

*NC* = Nature Credit

*NBI* = Net Biodiversity Impacts in quality hectare

*E*_t_ = Extent in hectares at project monitoring date

*C*_t_ = Condition at project monitoring date

*E_0_* = Extent in hectares at project start

*C_0_* = Condition at project start

*t* = Number of years from project start

*B* = Project crediting baseline

*L* = Leakage

Implementation in the LVMCM:

Calculation of the issuable Nature Credits for a given project can vary based on which metrics are selected as indicators. As the LVMCM grants the opportunity of observing the same site in different conditions, the reference site is defined as the project site in its undisturbed state immediately after assembly. The time variable has been kept constant, to allow for an unconfounded comparison with alternative credits. The following metrics were chosen as the three structure indicators: Abundance of the most abundant species, total biomass, and species richness. In the LVMCM, the abundance of the most abundant species is analogous to a metric such as canopy cover, as all species influence each other, and therefore the most abundant species has the most widespread impact. While total biomass and species richness provide direct information on the ecosystem’s structure, in accordance with Verra’s methodology. As for the two composition indicators, the LVMCM is not currently equipped to model between-guild competition. Therefore, instead of a species richness of a specific guild, selected indicator species’ abundances were used, in accordance with Verra’s methodology. Indicator species were assigned by sampling the species with the smallest negative impact on other species, and the species with the largest negative impact on other species. Further information on how the indicator species were chosen can be found below in section 2.4.1. The project crediting baseline (B) was kept constant at 0, to allow for an unconfounded comparison with alternative credits. This paper investigates biodiversity metrics and does not account for additional modifications to compute the final credits, such as accounting for leakage or a buffer. Therefore, NBI value was used in comparisons with other credits.

#### 5.2.6. The Wallacea Trust’s Biodiversity Credit

Wallacea trust define their Biodiversity Credit metric as a measured 1% uplift or avoided loss in biodiversity per hectare. Biodiversity values are measured as a median of a basket of minimum five metrics, one of which is an obligatory structural metric, while the rest are a score calculated from the relative abundance and conservation value of larger assemblages of species. Both the relative abundance and conservation values are simplified by thresholding into a five-point scale based on conformity with a pre-defined and analogous reference site. Subsequently, the biodiversity uplift is estimated based on the change in biodiversity values (Vm), and after accounting for a buffer and loss, the claimable Biodiversity Credits are issued.

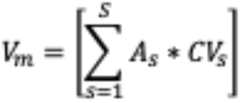

Where:

*Vm* = Overall biodiversity value for a sampling category

*As* = Relative abundance scores of each assigned on a 5-point scale

*CVs* = Conservation Value scores of each species is assigned on a 5-point scale

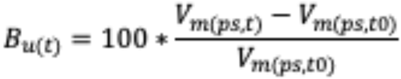

Where:

*Bu*(*t*) = Biodiversity uplift value for a given metric at a verification event

*Vm* = Overall biodiversity value for a sampling category

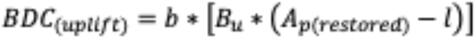

Where:

*BDC*(*uplift*) = Number of claimable biodiversity credits issued for an uplift project

*b* = Adjustment to account for the buffer retained by the registry issuing the credits for their insurance pool (this adjustment value is usually set at 0.8)

*Bu* = Median biodiversity uplift value across the different metrics after any adjustments have been made for uncertainty

*Ap*(*restored*) = Effective area in hectares within the Project Site restored over the Project

*l* = Number of hectares lost to leakage issues

Implementation in the LVMCM:

For the purposes of this research the focus is on biodiversity increase and not avoided loss, therefore we focus on Wallacea Trust’s Biodiversity Credit’s uplift calculations. As the LVMCM grants the opportunity of observing the same site in different conditions, the reference site is defined as the project site, in its undisturbed state immediately after assembly. For real-world ecosystems the Wallacea Trust’s methodology recommends three-dimensional data to be used as the obligatory structural metric. The LVMCM is a two-dimensional model, therefore the structural metric is interpreted as the overall biomass within the considered project area. Relative abundance of species is tracked throughout the experiments, while for the conservation value the IUCN Red List categories are used, in line with the Wallacea Trust’s methodology. Further information on how species are assigned IUCN Red List Categories can be found in the below in section 2.4.2. At any given time, a *Vm* value is calculated for all the species. Unfortunately, the LVMCM model in its current form only recreates intra-guild competition, therefore the requirement for measuring ‘larger assemblages of species’ is satisfied by grouping species based on their interspecies interaction characteristics. The chosen four groups of species are as follows: the group of species with the smallest negative impact on other species; the group of species with the largest negative impact on other species, the group of species with the largest overall impact on other species, and the group of species which is the most abundant in the reference condition, therefore having the most widespread impact on other species. Subsequently the change in the five metrics is calculated between degraded and restored states relative to the reference state. This paper investigates biodiversity metrics and does not account for additional modifications to compute the final credits, such as accounting for leakage or a buffer. Therefore, the *b* value was kept constant at 1.

### 5.3. Selection and implementation of biodiversity metrics

To help the contextualisation of the credits modelled, several established and independent biological metrics have also been implemented in the LVMCM. These are overall biomass, species richness, Range Size Rarity, Living Planet Index (Collen et al., 2009), Mean Species Abundance (Alkemade et al., 2009; Schipper et al., 2020), and the Species Threat Abatement and Restoration Metric (STAR) (Mair et al., 2021). Certain credits and metrics rely on locally or globally available data, such as indicator species, the IUCN Red List, and the IUCN Red List of Ecosystems (Keith et al., 2013; Myers et al., 2000; Rodrigues et al., 2006). A detailed description of the implementations of all is available below.

#### 5.3.1. Indicator Species

The LVMCM models complex community dynamics based on a few simple species characteristics: growth rate relative to the environment and species interactions. Species interactions are modelled through a network of randomly sampled adverse ecological interaction between the species. These interactions introduce an element of competition and thus limit the number of species that can co-exist at a given site. Accordingly, indicator species were chosen based on their relative impact on other species. The three indicator species are:

- The species that has adverse effects on most other species.
- The species that has adverse effects on least other species.
- The species with the largest relative impact on others. This species has the most detrimental effect on others, while being the least affected.

#### 5.3.2. IUCN Red List

The IUCN Red List is a worldwide database of species that categorises species based on their threat of extinction. The main categories are: Critically Endangered, Endangered, Vulnerable, Near Threatened, Least Concern. To apply these categories to the LVMCM’s simulated species, first the proportions of each category was noted according to the 2022-2 IUCN Red List, excluding species which are categorised as “Extinct”, “Extinct in Wild”, “Conservation Dependent” and “Data Deficient” (https://www.iucnredlist.org/statistics, EX=902; EW=84; CR=9251; EN=16364; VU=16493; CD=152; NT=8816; LC=77491; DD=20835; Total=128415). After the initial assembly of the community at each simulation, the simulated species were sorted into the IUCN Red List categories according to these proportions, based on their overall biomass. The highest biomass of each category was recorded as a threshold. After each further timestep simulated, for instance during a degradation or restoration event, the species were reorganised based these thresholds. Thus, if during a degradation experiment a “Vulnerable” species falls below the “Endangered” threshold, it is recategorized as “Endangered”.

#### 5.3.3. IUCN Red List Index (RLI)

The Red List Index (RLI) quantifies overall extinction risk for species based on their categorisation on the IUCN Red List. An RLI value of 1.0 indicates all species are classified as Least Concern, suggesting no imminent extinctions are expected. Conversely, an RLI of 0 signifies all species are considered extinct. The RLI for at a given time is calculated as:

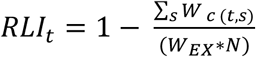

where:

*RLI_t_* = Red List Index at a given time

*W_c_* = IUCN Red List category weight of species s (Least Concern = 0; Near Threatened = 1; Vulnerable = 2; Endangered = 3; Critically Endangered = 4)

*t* = a given time

*s* = a given species

*W_EX_* = IUCN Red List category weight of for Extinct species = 5

*N* = Total number of assessed species

The LVMCM can track each species’ IUCN Red List category (see section 2.4.3), therefore the RLI could be straightforwardly implemented.

#### 5.3.4. IUCN Red List of Ecosystems

Similarly to the IUCN Red List of species, the IUCN Red List of Ecosystems were simulated based on the real-world proportions of each category: Critically Endangered, Endangered, Vulnerable, Near Threatened and of the Least Concern. (https://assessments.iucnrle.org/search statistics tab). Each site is categorised based on its overall Range Size Rarity index after the initial assembly of the metacommunity (See below). The thresholds for each category were noted. After each subsequent timestep modelled all sites were recategorized according to these thresholds. Therefore, if during a degradation experiment a “Vulnerable” ecosystem falls below the “Endangered” threshold, it is recategorized as “Endangered”.

#### 5.3.5. Living Planet Index

The Living Planet Index (LPI) measures the geometric mean of relative change in abundance of all vertebrate species compared to a baseline database from 1970. This was simulated as the geometric mean of the relative abundance of all species in the model. In the LVMCM, the initially assembled healthy community is chosen to be the baseline state. Any change from this state throughout subsequent experiment is measured, and the geometric mean is calculated using a regularisation constant chosen as the 1% of the monoculture carrying capacity of a single site.

#### 5.3.6. Mean Species Abundance

The Mean Species Abundance (MSA) aims to describe the ecosystem intactness and quantify ecosystem functioning. The MSA score is calculated by calculating the arithmetic mean for the change in species’ abundance between a project site and an undisturbed reference site (Alkemade et al., 2009; Schipper et al., 2020). To calculate the change in species’ abundance the abundance at the project site is divided by the abundance at the reference site, while truncating the values at 1. Importantly, invasive species at the project site which are not present at the reference site are not counted. Again, in the LVMCM the initially assembled healthy community is used as the reference state. MSA is computed on a site-by-site basis which then is averaged to calculate the MSA for a given area in question.

#### 5.3.7. Range Size Rarity

Range Size Rarity indicates the relative conservation importance of a given location. For a given species the Range Size Rarity of a given location is the proportion of a species’ range contained within the location. By summing the Range Size Rarity values for all species at a given location the overall importance of the location for conservation can be measured. Range Size Rarity is implemented on a site-by-site basis in the LVMCM in a straightforward manner. Presence or absence at a site was determined by the simulation’s extinction threshold.

#### 5.3.8. Species Threat Abatement and Restoration Metric

The Species Threat Abatement and Restoration Metric (STAR) measures the impact of changes in an ecological community on species extinction risk (Mair et al., 2021). There are two STAR metric values: The STARt score shows the level of threat facing a given species at a given location, while the STARr score shows the effort required to get a given species to the IUCN Red List’s Least Concern category. Thus, the sum of STAR scores for all species at a given location represent a proportion of the global opportunity to reduce species’ extinction risk through threat abatement and restoration, respectively. The sum of STAR scores for a given species over all locations represents the global effort needed to get a species to the IUCN Red List’s Least Concern category. The STARt metric is calculated as:

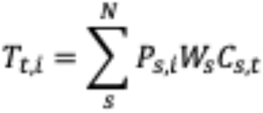

Where:

*T_t,i_* = Threat Abatement metric

*P_s,_*_I_ = the current area of habitat of each species s within location i (expressed as a percentage of the global species’ current area of habitat)

*W_s_* = IUCN Red List category weight of species s (Near Threatened = 1; Vulnerable = 2; Endangered = 3; Critically Endangered = 4)

*C_s,t_* = The relative contribution of threat t to the extinction risk of species s

*N* = The total number of species at location i

In the LVMCM, the STARt score is straightforward to implement based on the simulated IUCN Red List categories and Range Size Rarity calculations. The Area of Habitat for a given species was determined based on local presence – absence data from the pristine state. Importantly, as the model does not distinguish between differing threats, C is kept constant at 1. This paper investigates biodiversity metrics and does not account for additional modifications, such as accounting for leakage or buffers. Therefore, the STARr score is not considered, as the habitat specific restoration multiplier is not possible to compute.

