## Supplementary figures and images for "Evaluating Biodiversity Credit Metrics Using Metacommunity Modelling"

### Figure 1.png

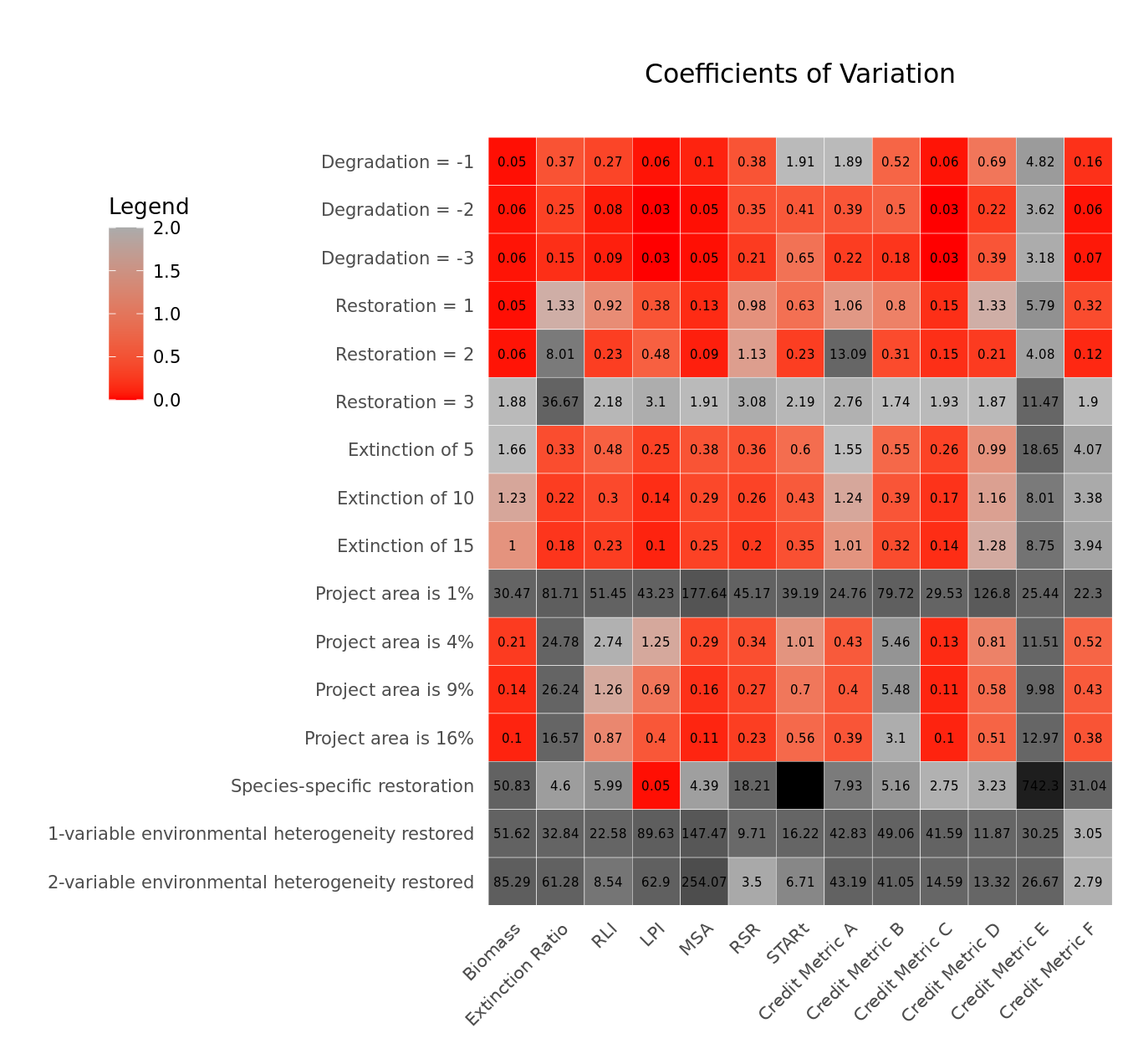

### Figure 2.png

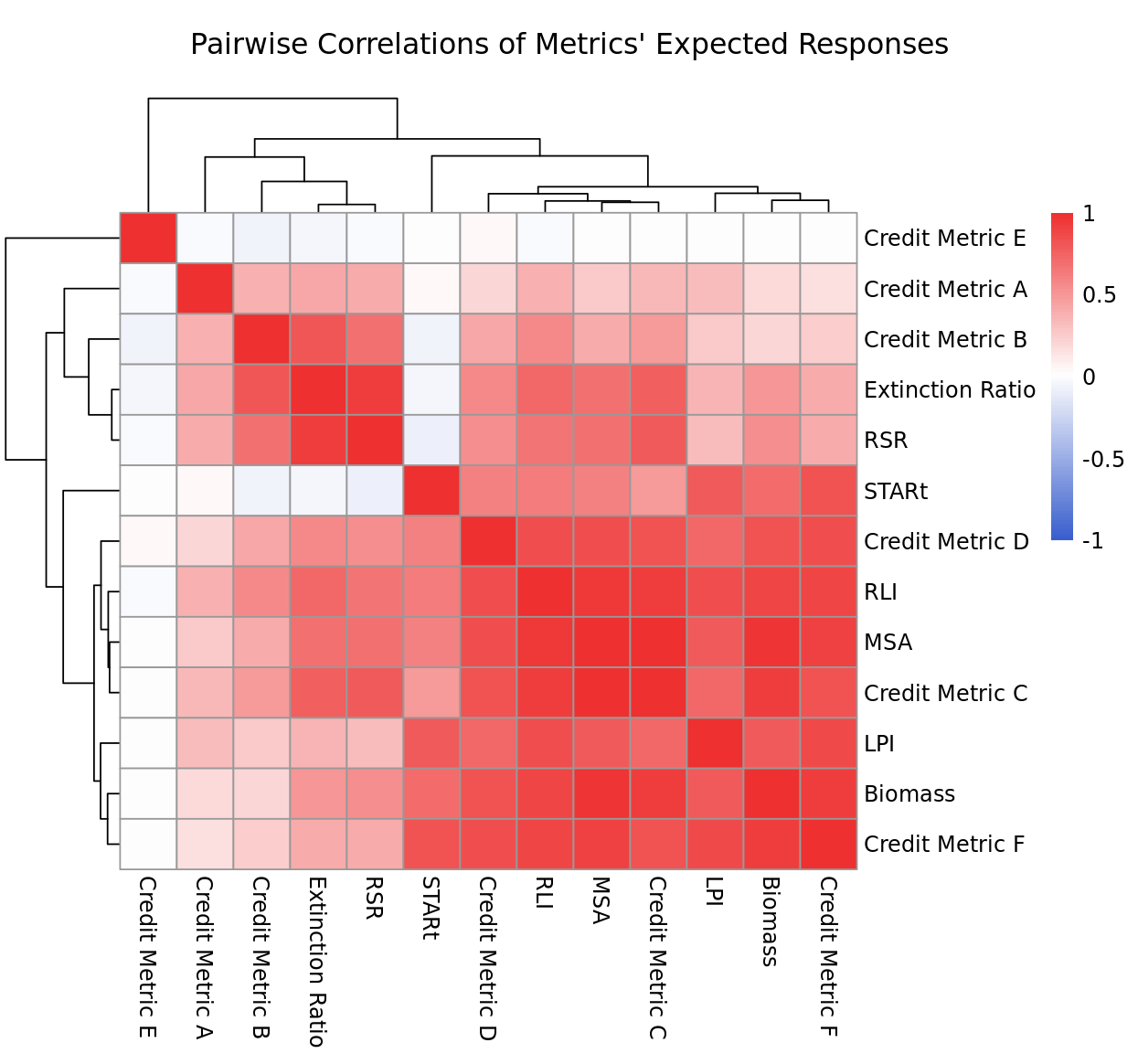

### Figure 3.png

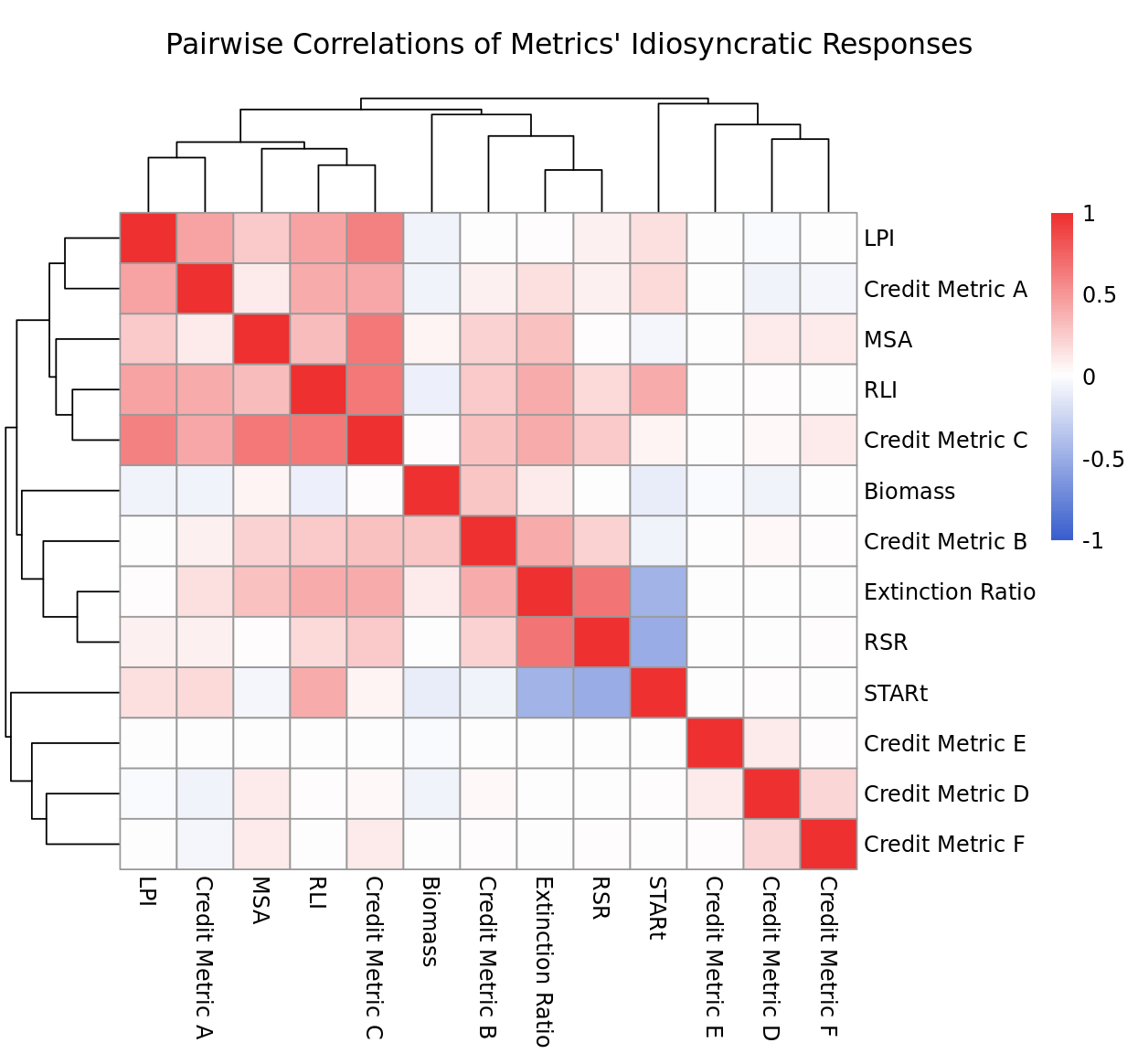

### Figure 4.png

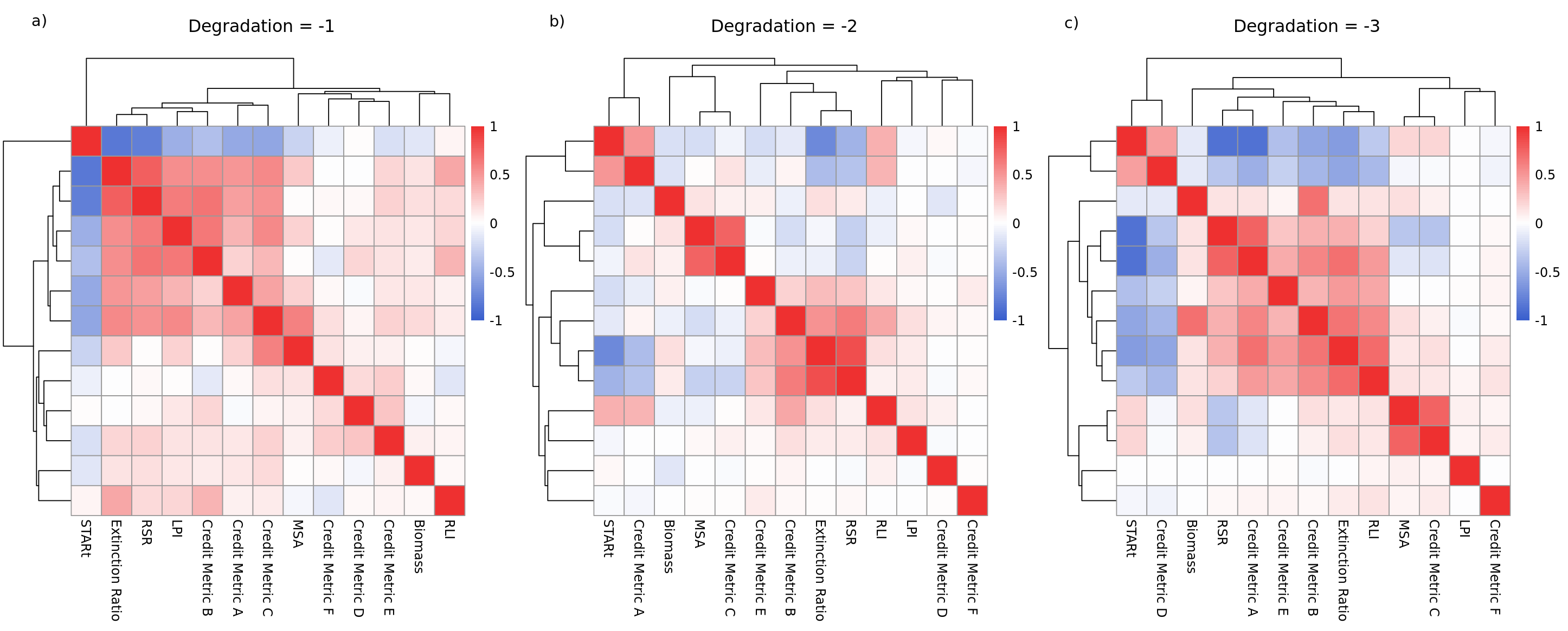

### Figure 5.png

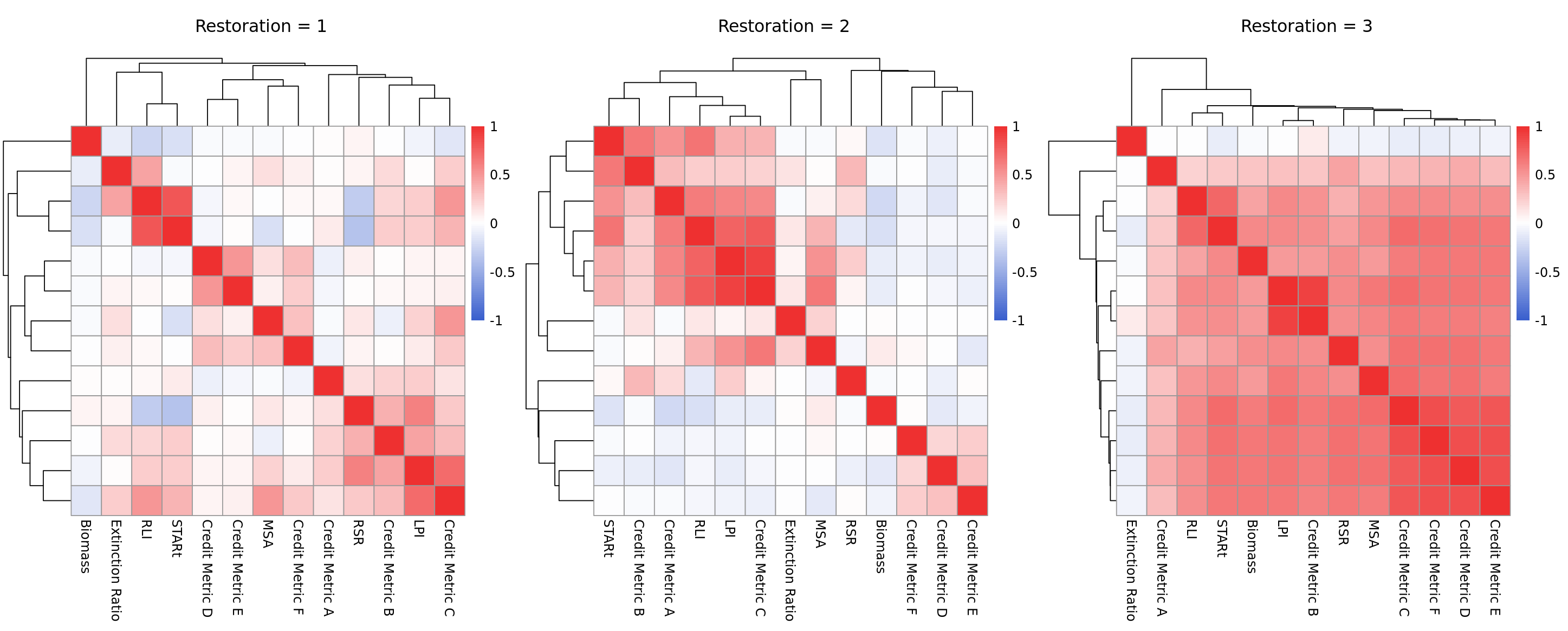

### Figure 6.png

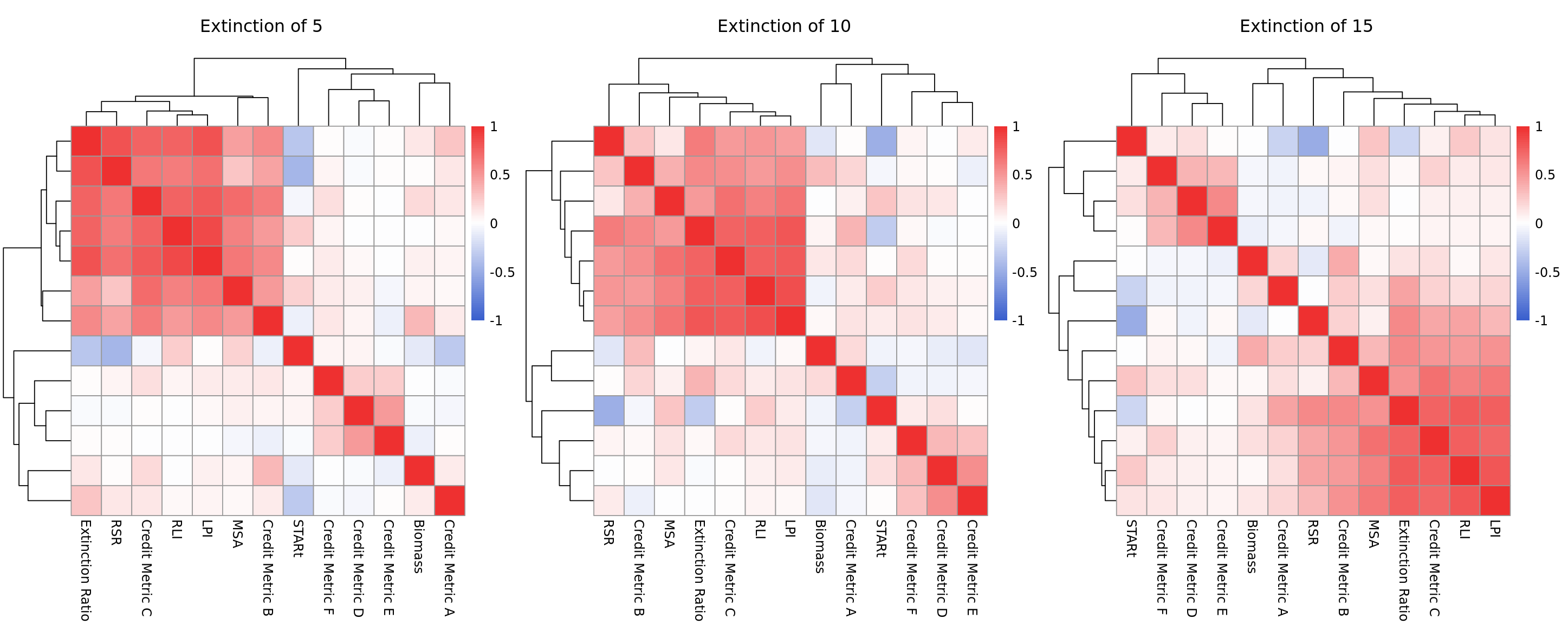

### Figure 7.png

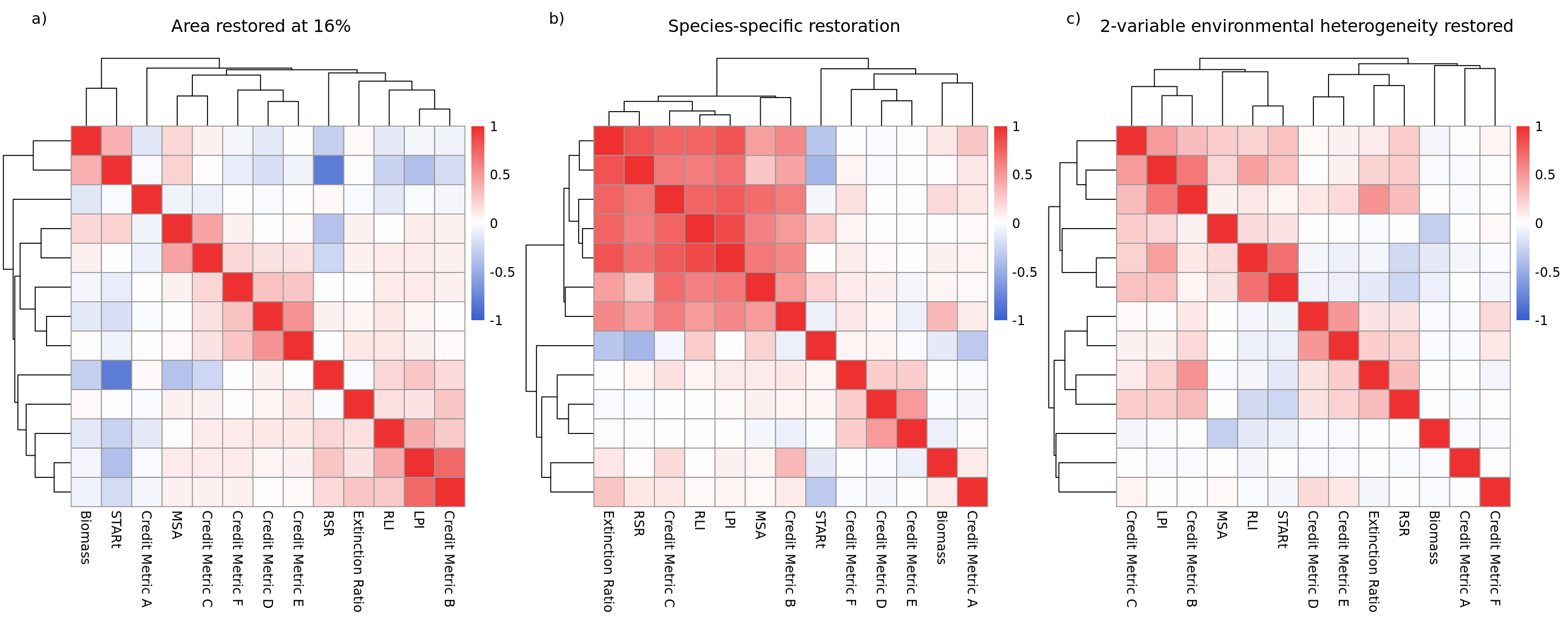

### Saturated ecosystem.png

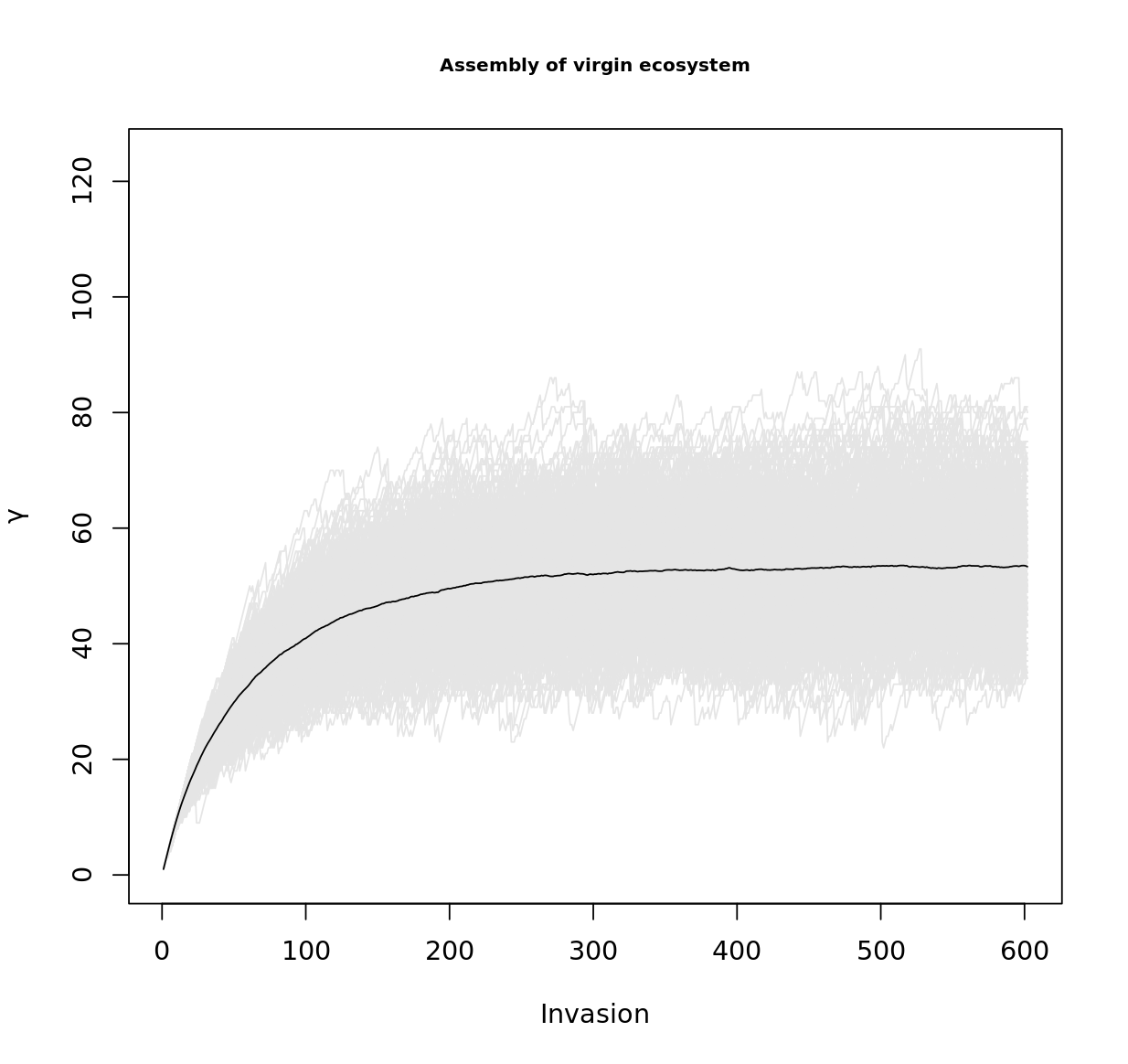
